# Respiratory viral infections prime accelerated lung cancer growth

**DOI:** 10.1101/2025.09.02.672566

**Authors:** Wei Qian, Xiaoqin Wei, Andrew J Barros, Xiangyu Ye, Qing Yu, Samuel P Young, Eric V Yeatts, Yury Park, Chaofan Li, Gislane Almeida-Santos, Jinyi Tang, Harish Narasimhan, Nicole A Kirk, Ying Li, Li Li, Peter Chen, Jeffrey M Sturek, Kwon-Sik Park, Wei Chen, In Su Cheon, Jie Sun

**Author notes:** These authors contributed equally (Wei Qian, Xiaoqin Wei).

## Abstract

The COVID-19 pandemic has highlighted long-term health concerns of viral pneumonia, yet its potential impact on cancer development and growth remains poorly understood. Here, we demonstrate that prior infection with SARS-CoV-2 or influenza virus promoted lung tumor progression by reprogramming the local immune landscape. Retrospective clinical analysis revealed that patients hospitalized with COVID-19 exhibited increased lung cancer incidence. Using multiple murine lung cancer models, we show that prior severe respiratory viral infections accelerated tumor growth and reduced survival. Mechanistically, prior viral pneumonia epigenetically remodeled the lung to establish a pro-tumor microenvironment, including the local accumulation of SiglecF^hi^ tumor-associated neutrophils, a transcriptionally reprogrammed, immunosuppressive population whose signature predicted poor prognosis in human lung adenocarcinoma. In parallel, epithelial compartments exhibited altered differentiation trajectories, with persistence of injury-associated alveolar intermediates positioned along tumorigenic lineages. We observe sustained chromatin remodeling at key cytokine loci in immune and structure cells, linking inflammatory memory to persistent immune suppression. Therapeutically, combined inhibition of neutrophil recruitment via CXCR2 and PD-L1 signaling restored CD8⁺ T cell infiltration and suppressed tumor growth. Together, our findings establish a direct causal relationship between viral pneumonia, including COVID-19, and lung tumorigenesis, highlighting the urgent need to monitor survivors for elevated cancer risk and to develop targeted interventions and therapies aimed at preventing potential cancer bursts in COVID-19 convalescents.

**Graphic Abstract:** 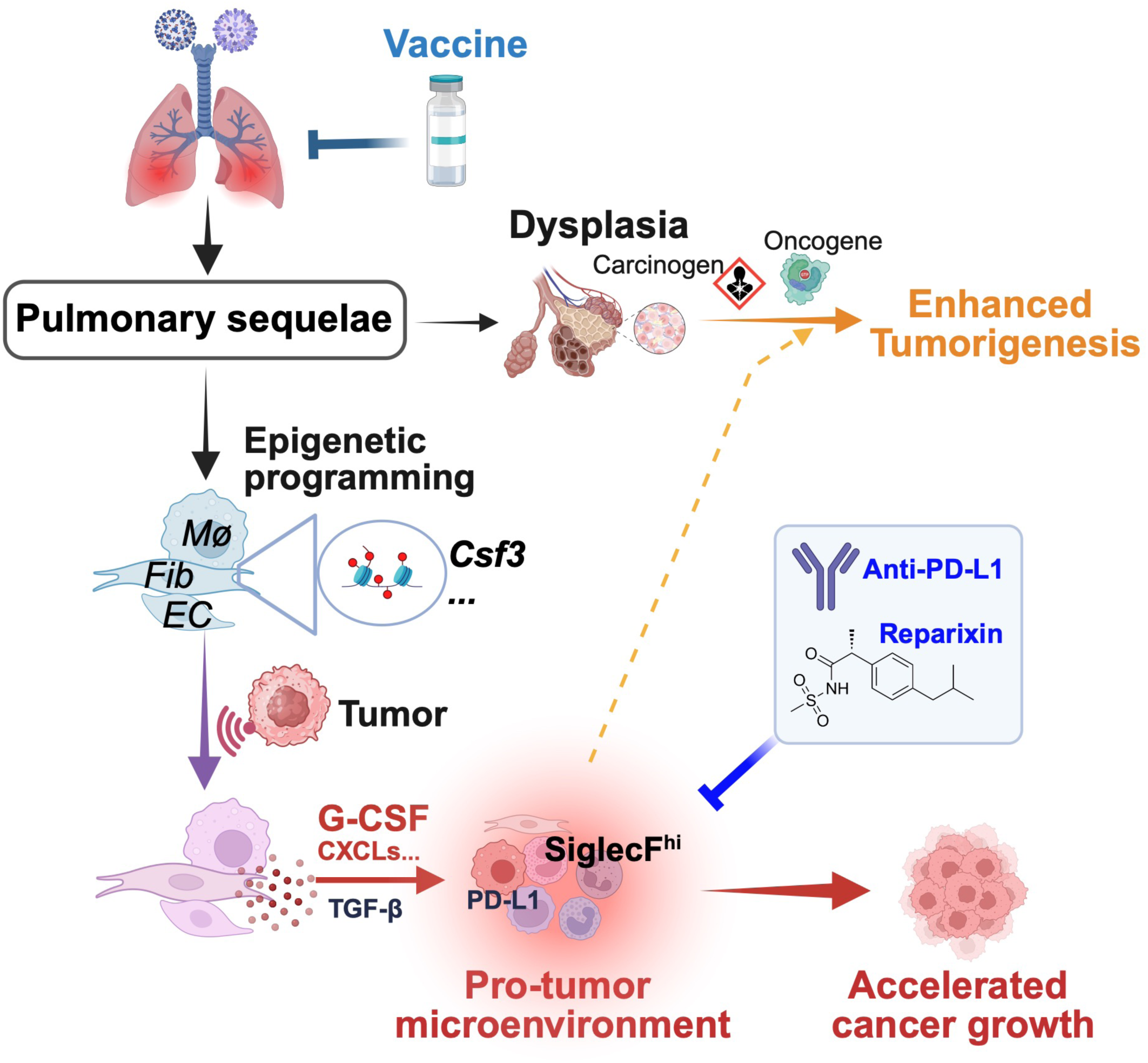

## INTRODUCTION

The COVID-19 pandemic has fundamentally reshaped global health, revealing infection-induced long-term complications that persist beyond the resolution of acute illness.^1^ As the world transitions into the post-pandemic era, growing evidence highlights persistent complications among COVID-19 patients, referred to as post-acute sequelae of SARS-CoV-2 (PASC, also known as long COVID). While most individuals recover within weeks, a substantial number continue to experience lingering symptoms, systemic dysfunction, and chronic inflammation.^2–4^ Beyond COVID-19, other respiratory viral infections, notably influenza, have also been associated with prolonged health issues^5,6^, sometime termed as “long flu”.^7^ Though underexplored, data have suggested that influenza infection may predispose individuals to increased lung cancer risks.^8^ Given the comparatively recent onset of the COVID-19 pandemic, robust epidemiological data linking COVID-19 pneumonia to lung cancer are still limited. However, a small cohort study indicates that individuals previously infected with SARS-CoV-2 may face an elevated risk of respiratory diseases, including lung cancer.^9^

Chronic inflammation is widely recognized as a key driver of cancer development, including lung tumorigenesis.^10,11^ COVID-19, particularly PASC, elicits persistent immune activation characterized by the overproduction of inflammatory cytokines such as IL-6, IL-1β, and TNF.^12–14^ This sustained inflammatory milieu may cause oxidative stress, DNA damage and genomic instability to potentially facilitate cell transformation.^15^ Persistent immune dysregulation in post-viral syndromes may also compromise immune surveillance, allowing pre-malignant cells to escape recognition and evolve into malignant phenotypes.^16^ Furthermore, inflammation-driven changes in the lung microenvironment may create favorable systemic or local conditions for tumor progression.^17^ Despite these insights, a direct causal link between prior viral pneumonia with later cancer development remain unestablished. Given the vast number of individuals globally affected by SARS-CoV-2 and influenza, and the growing recognition of long-term sequelae such as pulmonary fibrosis and immune dysfunction, clarifying how respiratory viruses may influence cancer risk is an urgent priority. If such infections indeed promote tumorigenesis or cancer growth, identifying the underlying molecular and immunological mechanisms will be crucial for establishing early detection strategies, preventive interventions and targeted therapies.

Neutrophils are increasingly recognized for their multifaceted roles in cancer. Tumor-associated neutrophils (TANs) can exhibit both tumor-promoting and tumor-restraining activities, depending on the context, tumor stage and local cytokine cues.^18–20^ Emerging single-cell and spatial profiling studies have revealed distinct TAN subsets with roles in immunosuppression, angiogenesis and matrix remodeling, but the mechanisms that drive their differentiation and persistence in tumors are not fully understood.^21–24^ In lung adenocarcinoma (LUAD), the presence of neutrophil-rich microenvironments has been associated with poor prognosis and resistance to immune checkpoint blockade,^25,26^ yet it remains unclear how these cells are recruited, programmed, or maintained *in situ*.

Moreover, the potential contribution of prior inflammatory or infectious insults to TAN expansion and functional reprogramming has not been explored. Addressing these questions is important for targeting specific TAN phenotypes as therapeutics or biomarkers of tumor progression and immune evasion.

Here, we investigate whether and how prior severe respiratory viral infections shape the lung environment to promote tumorigenesis. Combining retrospective clinical analysis in a large clinical database with mechanistic studies in multiple murine lung cancer models, we define the long-term lung remodeling process following viral pneumonia including SARS-CoV-2 and influenza that accelerates tumorigenesis and cancer growth. Using single-cell transcriptomic and epigenomic profiling, functional immune analyses and therapeutic interventions, we revealed that persistent epigenetic changes in inflammatory cytokine loci after infection established a lung microenvironment prone to cancer, driven in large part by elevated pro-tumor neutrophil activity *in situ.* Together, these findings provide a mechanistic framework linking severe viral pneumonia to increased lung cancer risk, and highlight potential pathways for early screening and intervention.

## Results

### Prior severe COVID-19 is associated with increased lung cancer risk

To investigate the relationship between SARS-CoV-2 infection and lung cancer incidence, we conducted a retrospective cohort study using Cosmos Epic database. A total of 44,229,908 (aged 55-74 years) non-infected control, non-hospitalized (ambulatory) or hospitalized COVID-19 patients were analyzed in the first two years of COVID-19 pandemic (2020-2021), with baseline demographic characteristics shown in Table S1. Cancer diagnosis was evaluated across varying severities of COVID-19 infection from January 2022 onwards. Our analysis revealed that patients who experienced severe COVID-19, defined by hospitalization, had a modest elevated risk of developing all cancer types overall, but diminished incidence in male prostate cancer and female breast cancer. Notably, while prior non-severe SARS-CoV-2 infection is associated with a moderate decrease in lung cancer diagnosis, prior hospitalized COVID-19 resulted in a 1.19-fold higher hazard ratio (HR) for lung cancer diagnosis compared to uninfected controls after adjustment with gender, age and smoking status (Figure 1A). We further stratified cumulative hazard by COVID-19 severity and smoking history. Across current, former, and never smokers/unknown, patients who were hospitalized with COVID-19 consistently demonstrated the highest cumulative hazard for lung cancer diagnosis compared to both ambulatory and uninfected controls (Figure 1B). Similar increased risks of lung cancer incidence have also been observed after influenza infection,^8^ indicating that increased risk of lung cancer development and/or growth likely represent a common theme after respiratory viral infections.

**Figure 1.**
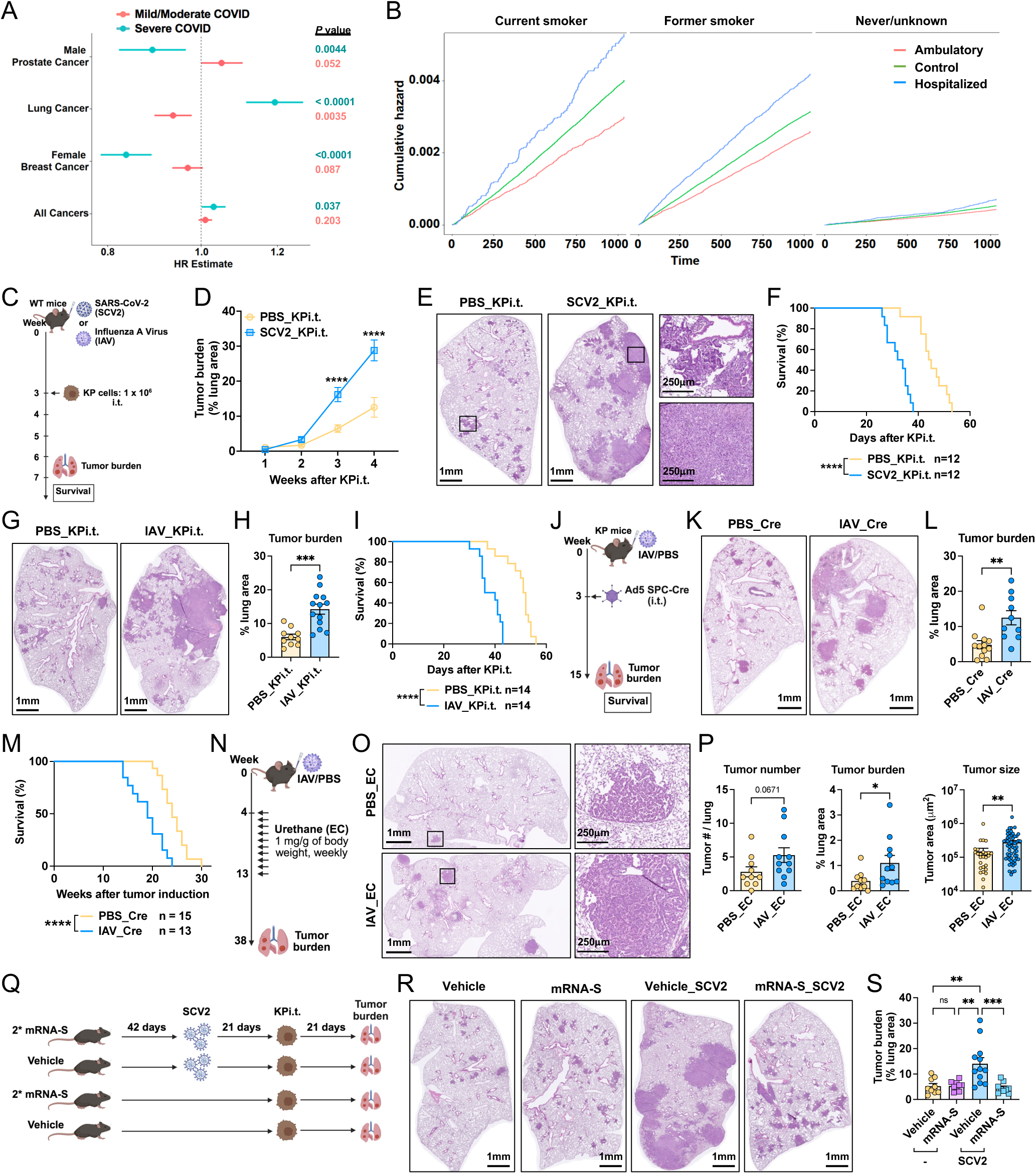
Acute respiratory viral infections prime enhanced lung tumor growth, which is prevented by vaccination. (**A**) Analysis of the Cosmos database comparing lung cancer risk (hazard ratios) in individuals with (severe) or without (mild/moderate) prior COVID-19 hospitalization. (**B**) Cumulative incidence of lung cancer stratified by COVID-19 severity and smoking status. (**C-I**) Experimental model evaluating the impact of prior respiratory viral infection on tumor growth and survival. (C) Schematic of intratracheal (i.t.) KP tumor cell transfer at day 21 post SARS-CoV-2 (SCV2) or influenza A virus (IAV) infection, followed by tumor burden and survival assessments. (D) Tumor growth kinetics from week 1 to week 4 after KP cell inoculation in SCV2-versus PBS-treated mice. (E) Representative tumor-bearing lung lobe at week 3. (F) Survival analysis of SCV2-infected and uninfected mice following tumor inoculation. Representative lung lobe (G) and tumor burden quantification (H) at week 3 in IAV- and PBS-treated groups. (I) Survival analysis of IAV-infected and control mice post tumor implantation. (**J-M**) Prior IAV infection promoted lung tumor development in a genetically engineered mouse model, Kras^LSL-G12D^; p53^fl/fl^ (KP) mice. (J) Schematic of experimental design: KP mice were inoculated with PBS or IAV and, three weeks later, received i.t. Ad5-SPC-Cre virus to initiate tumor development. Representative lung lobe (K) and tumor burden quantification (L) at 12 weeks post-tumor initiation in IAV-versus PBS-treated mice. (M) Survival analysis of tumor-bearing KP mice with or without prior IAV infection. (**N-P**) Prior IAV infection accelerated tumorigenesis in a chemical-induced lung cancer model. (N) Experimental setup. Mice were infected with IAV for four weeks before received weekly intraperitoneal injections of urethane (ethyl carbamate, EC) for 10 weeks. Tumor burden was assessed at week 34 (8 months). (O) Representative tumor H&E images. (P) Quantification of tumor burden, tumor number, and size in PBS_EC and IAV_EC mice. (**Q-S**) SARS-CoV-2 spike mRNA vaccination mitigated infection-induced tumor promotion. (Q) Schematic of mRNA vaccination protocol and tumor challenge. Mice were immunized with vehicle or SARS-CoV-2 spike mRNA (2 doses, 21-day interval), followed by SCV2 or PBS infection. KP tumor cells were inoculated 21 days post-infection, and tumor burden was assessed after another 21 days. (R) Representative histological lung sections. (S) Quantification of tumor burden in each group. Data represent at least two independent experiments or pooled from two (D, P, and S) and three (F, H, I, L, and M) experiments. Graphs display mean ± SEM. Statistical significance was assessed by two-way ANOVA (D), Mann-Whitney test (L and P), one-way ANOVA (S), and Log-rank Mantel-Cox test (F, I and M). (ns *p* > 0.05, \**p* < 0.05, \*\**p* < 0.01, \*\*\**p* < 0.001, \*\*\*\**p* < 0.0001)

### Acute respiratory viral infections prime increased lung tumor development

Next, we sought to establish animal models to mechanistically investigate the link of prior respiratory viral infection with lung cancer development and/or growth. We first employed an orthotopic mouse model in which Kras^G12D^ and p53-deficient (KP) lung tumor cells were intratracheally (i.t.) delivered into mouse (Figures S1A-S1C). To do this, 6-month-old wild-type C57BL/6 male mice were intranasally infected with the mouse adapted SARS-CoV-2 (SCV2) strain MA10. Middle-aged male mice were used as these mice were shown to be more susceptible to SARS-CoV-2 infection and diseases.^27,28^ At 21 day post-infection (d.p.i.) when virus has been cleared from the host for about two weeks,^27,29^ mice were received KP tumor cells and tumor burden or survival were monitored (Figure 1C). Prior SARS-CoV-2 infection had no apparent impact on initial tumor development, as evidenced by similarly low tumor burden and sparse, localized lesions in both mock-infected (PBS) or SARS-CoV-2-infected groups at week 1. By week 2, tumors in the infected group began to show increased growth relative to PBS controls, although differences in overall burden remained non-prominent at this stage. However, from week 3 onward, infection-experienced mice exhibited pronounced tumor expansion, forming large, confluent masses occupying substantial portions of the parenchyma. In contrast, PBS-treated mice maintained relatively preserved alveolar architecture, characterized by scattered and small tumor nodules (Figures 1D, 1E, and S1D). Consistent with this increased tumor burden, infection-experienced mice demonstrated significantly reduced survival relative to uninfected controls (Figure 1F). Since prior clinical study has also linked influenza infection to an increased lung cancer incidence in humans,^8^ we sought to determine whether this tumor-promoting effect observed with SARS-CoV-2 infection was also applicable to influenza virus A/PR8/34 (IAV) infection. To this end, we infected mice with IAV and subsequently challenged them with KP tumor cells after viral clearance (typically at 10 d.p.i.^30^) at 21 d.p.i. (Figure 1C). Similar to what was observed from SARS-CoV-2 infection, prior IAV-infected mice displayed significantly increased tumor burden and mortality compared to PBS-treated controls (Figures 1G-1I). Notably, this pro-tumorigenic effect persisted even when KP cells were inoculated 60 days post-IAV infection (Figure S1E). Comparable phenotype was also observed using Lewis lung carcinoma (LLC) cells (Figure S1F), indicating that the phenomenon is not limited to a single tumor cell type.

Compared to those of SARS-CoV-2 infection in mice, mouse IAV infection generally induced higher levels of lung damage and more prolonged tissue inflammation, which appeared to better mimics persistent lung sequelae observed after acute SARS-CoV-2 infection in humans as shown previously.^27,31^ Therefore, we sought to use the model to examine whether prior viral pneumonia could promote lung cancer development in a genetically engineered mouse model (GEMM) of lung cancer, which take substantially longer time for lung cancer development.^32^ To this end, we utilized the induced Kras G12D expression and p53 gene loss (KP) mice to assess whether prior viral pneumonia accelerates lung tumor progression. In prior IAV-infected mice, intratracheal inoculation with Ad5-SPC-Cre to initiate lung tumorigenesis resulted in significantly greater tumor growth at 12 weeks after tumor induction and reduced survival compared to uninfected controls (Figures 1J-1M). Additionally, in a chemically induced lung cancer model using urethane (ethyl carbamate, EC),^33^ prior severe IAV infection exacerbated tumorigenesis, leading to increased tumor number, area, and size at eight months post-treatment (Figures 1N-1P). Collectively, these data provide a direct experimental evidence that severe respiratory viral infections prime the lung for enhanced tumorigenesis and cancer progression.

### Prior mRNA vaccination mitigates enhanced cancer growth after viral infection

Vaccination is the most effective approaches of preventing acute and chronic viral-associated diseases.^34^ To determine whether mRNA vaccination could mitigate infection-primed tumor progression, mice were intramuscularly immunized with two doses of mRNA-Spike (mRNA-S) vaccine or vehicle control at a 21-day interval. Mice were then infected with SARS-CoV-2 and received KP tumor cells at 21 d.p.i. Tumor burden was assessed three weeks later (Figure 1Q). Two doses of mRNA-S provided full protection against mouse morbidity after infection (Figures S1G and S1H).

Importantly, mRNA-S immunized mice exhibited significantly lower tumor burden compared to mock immunized mice after infection, with tumor levels comparable to uninfected controls (Figures 1R and 1S). Furthermore, mRNA-S vaccination alone did not increase tumor burden compared to the vehicle control. These findings suggest that mRNA vaccination protects against lung tumor development, likely by preventing severe disease development after viral exposure. Supporting this notion, prior mild infections with low dose SARS-CoV-2 or IAV did not significantly increase tumor burden compared to uninfected controls, whereas severe infections did (Figures S1I-S1L). These results highlight the critical role of infection severity in promoting lung tumor growth, which is supported by the clinical observations above (Figures 1A and 1B).

### Prior respiratory viral infections program a pro-tumor local environment

To elucidate the mechanisms underlying infection-induced tumor promotion, we performed single-cell RNA sequencing (scRNA-Seq) on whole lung cells from KP mice at 6 (PBS_T6 and IAV_T6) and 12 (PBS_T12 and IAV_T12) weeks post-tumor induction, as well as from tumor-free controls (PBS and IAV, time point equivalent to T6) (Figure 2A). Unbiased clustering of all cells revealed a diverse array of cell populations, including different immune cells, endothelial cells, fibroblasts, normal and transformed epithelial cells (Figure S2A).

**Figure 2.**
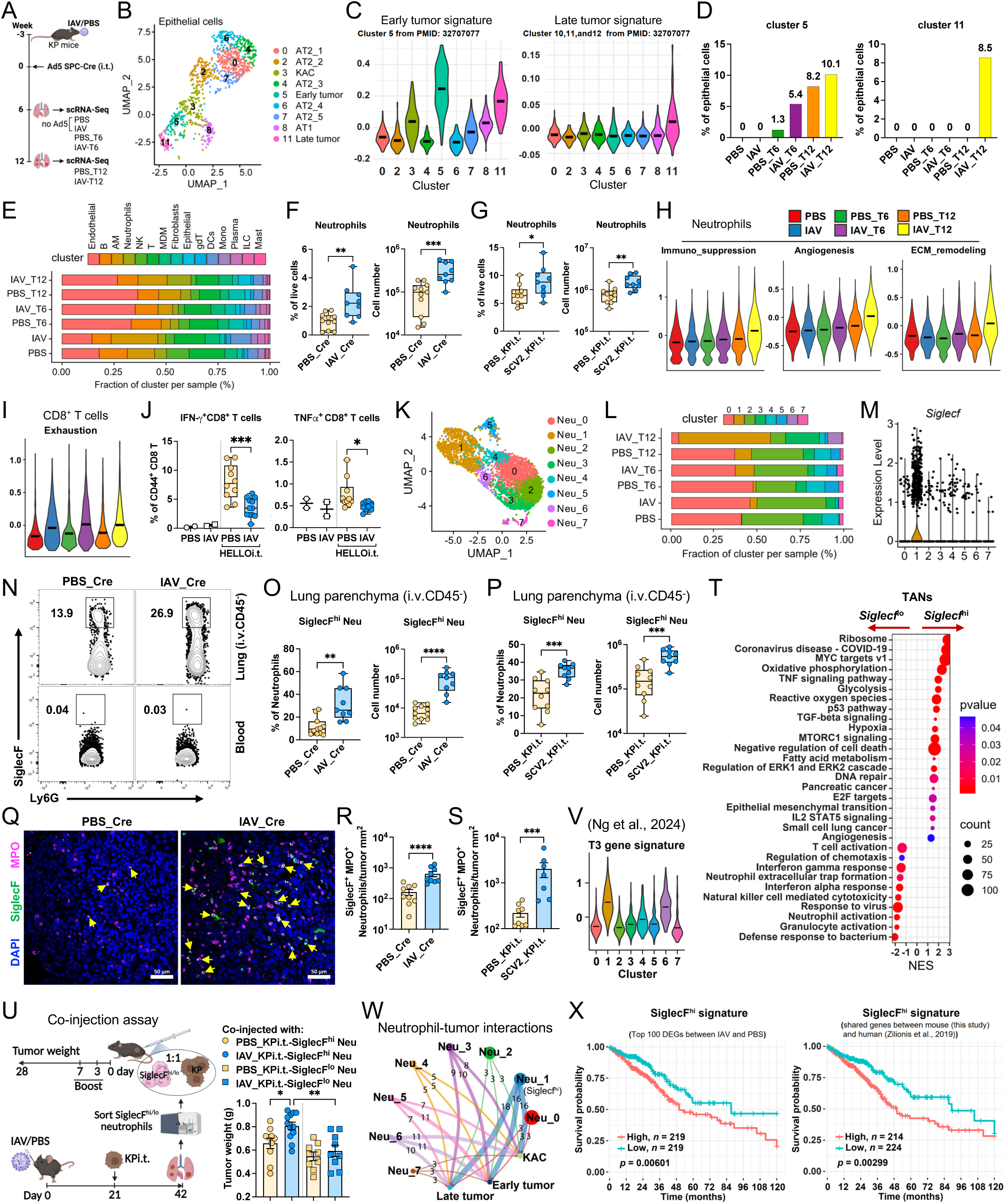
Prior respiratory viral infections program pro-tumor local environment. (A) Schematic showing scRNA-Seq experimental setup. KP mice were infected with IAV for 3 weeks, followed by i.t. administration of Ad5 SPC-Cre virus to initiate tumorigenesis. Lung single-cell suspensions were subjected to scRNA-Seq at weeks 6 (early-stage) and 12 (late-stage) after tumor induction. Tumor free PBS and IAV groups served as controls. Each group was representing pooled samples from 3-5 mice. (B) UMAP projection of alveolar epithelial cell populations including AT2, AT1, and KRT8+ alveolar intermediate cells (KAC), and clusters corresponding to early and late tumor cells. (C) Violin plots showing early tumor (cluster 5, left) and late tumor (cluster 11, right) signature scores across the 9 alveolar epithelial clusters identified in (B). (D) Proportions of clusters 5 and 11 among total epithelial cells. (E) Distribution of major clusters of immune cells per group. (**F and G**) Frequency and absolute number of total neutrophils in the lung parenchyma (i.v.CD45^-^) of tumor-bearing KP mice with prior IAV or PBS infection (F), and in SCV2_KPi.t. and PBS_KPi.t. mice (G). (H) Neutrophil pro-tumor signature scores across experimental groups. (I) CD8^+^ T cell exhaustion signature scores across the same groups as in (H). (J) Percentage of IFN-γ^+^ or TNFα^+^ CD8^+^ T cells among CD44^+^ CD8^+^ T cells in tumors of IAV- or PBS-pre-exposed mice, 3 weeks post KP-HELLOi.t. tumor implantation. (**K and L**) UMAP projection (K) and proportion (L) of identified neutrophil subsets. (**M**) Violin plot displaying *Siglecf* expression levels among neutrophil clusters. (**N**) Flow cytometry dot plots showing the presence of SiglecF^hi^ neutrophils in tumor-bearing lungs and blood of PBS-Cre or IAV-Cre mice. (**O and P**) Frequency and absolute number of SiglecF^hi^ neutrophils among total tumor-infiltrating neutrophils in PBS_treated versus IAV_Cre (O) or SCV2_KPi.t. (P) mice. (**Q**) Representative images of immunofluorescence staining showing co-localization of MPO and SiglecF in tumors from PBS_Cre and IAV_Cre. Scale bar, 50 μm. (**R and S**) Quantification of SiglecF^+^ MPO^+^ neutrophils per mm^2^ tumor area in IAV_Cre (R) and SCV2_KPi.t. (S) tumor sections. (T) Pathways enrichment analysis (Hallmark and GO database) of DEGs between *Siglecf*^hi^ and *Siglecf^l^*° neutrophils. Cutoffs: p.adj < 0.05; NES > 1. (U) Schematic of neutrophil-KP tumor cell co-injection experiment. Equal numbers of sorted neutrophils and KP tumor cells were co-injected s.c. into the right flank of naïve recipient mice. Neutrophils were further boosted at the tumor site via s.c. injection on days 3 and 7. Tumor growth and endpoint weight (day 28) were assessed. Right: tumor weights following co-injection of PR8- or PBS-conditioned SiglecF^hi^ or SiglecF^lo^ neutrophils. (V) Violin plot showing reported pro-tumor neutrophil (T3) gene signature scores across neutrophil clusters identified in (K). (W) CellChat analysis of ligand-receptor interactions between neutrophil clusters and tumor cell subtypes (KAC, early tumor, late tumor). (X) Overall survival analysis of lung adenocarcinoma patients in TCGA stratified by gene signature of SiglecF^hi^ neutrophils from either top 100 DEGs between IAV-SiglecF^hi^ TANs and PBS-SiglecF^hi^ TANs, or shared genes between mouse and human. Data represent two (Q) or three (N) experiments, and pooled from two (G, J, P, R, and S) or three (F, O, and U) experiments. Graphs display mean ± SEM. Statistical significance was assessed by Mann-Whitney test (F, G, J, O, P, R, and S) and one-way ANOVA (U). (\**p* < 0.05, \*\**p* < 0.01, \*\*\**p* < 0.001, \*\*\*\**p* < 0.0001)

In order to understand how infection alters epithelial cell states in the context of tumorigenesis, we interrogated the transcriptional landscape of the epithelial compartments. Subclustering of epithelial cells identified two major lineages: airway and alveolar epithelial cells. The airway lineage included basal, ciliated and club cell populations (Figures S2B-S2E), while the alveolar lineage comprised AT2 cells (c0, c2, c4, c6, c7), AT1 cells (c8), and KRT8^hi^ alveolar intermediated cells (KACs, c3). Clusters 5 and 11, which expressed *Krt8*, also emerged as transcriptionally distinct epithelial subsets, suggesting potential state transitions within the alveolar lineage (Figure 2B). To better understand the identity of these epithelial subpopulations, we assessed their functional states using cell identity scoring. We found c5 and c11 exhibited the lowest AT2-like scores, indicating the loss of AT2 identity, while KACs (c3) displayed elevated AT1-like scores, suggesting a transitional state (Figure S2F; Table S2). Pseudotime and CytoTRACE analyses revealed that KACs occupies a differentiation trajectory leading toward either AT1 cells or more transformed cell states, consistent with prior reports implicating KRT8^hi^ intermediates serve as progenitors for LUAD development (Figure S2G).^35^ To evaluate whether c5 and c11 represented tumor cell states, we applied previously defined early and late tumor gene signatures,^36^ and found that c5 exhibited highest early tumor signature, while c11 corresponded to late-stage tumors, indicating a progression axis with the epithelial compartment (Figure 2C). Importantly, IAV-experienced mice had an increased proportion of early tumor c5 cells at both early (T6) and late (T12) time points (Figure 2D). Remarkedly, the late tumor c11 was exclusively present in IAV_T12 samples (Figures 2D, S2D and S2E), implying that prior IAV infection promotes tumor progression.

Cancer growth is profoundly shaped by the surrounding immune microenvironment.^37^ To determine whether prior IAV infection changes the tumor microenvironment (TME), we examined immune cell composition and function across experimental groups. Notably, we observed significant alterations in cell composition, particularly a marked expansion in neutrophils in IAV_T12 tumors compared to all other groups (Figure 2E). This increase was further confirmed by flow cytometry in both IAV GEMM and SARS-CoV-2 KPi.t. tumor transfer models, where lung parenchymal neutrophils (i.v.CD45^-^), but not blood neutrophils, were higher in prior infected tumor-bearing mice (Figures 2F, 2G, and S3A-S3C). Beyond cell abundance, neutrophils from IAV-associated tumors displayed higher pro-tumor signatures than those in PBS-treated counterparts at both early and late tumor stages (Figures 2H and S2H). Conversely, anti-tumor neutrophil signatures were downregulated in the IAV groups (Figure S2I; Table S2), indicating a functional reprogramming toward a tumor-promoting phenotype. In parallel, alveolar macrophages (AMs) and interstitial macrophages (IMs) from IAV_T12 lungs also showed enhanced pro-tumor signatures (Figure S2J; Table S2). Meanwhile, CD8^+^ T cells exhibited increased exhaustion and reduced effector function gene programs in the context of prior infection (Figures 2I and S2K; Table S2). To functionally validate the impact of prior infection on anti-tumor T cell responses, we employed KP tumor cells expressing CD4 and CD8 epitopes from lymphocytic choriomeningitis virus (LCMV) glycoprotein (KP-HELLO).^38^ When these tumor cells were i.t. transplanted into IAV-infected or control mice, and subsequently restimulated the lung cells with LCMV GP33 peptide, CD8^+^ T cells from prior infected mice exhibited markedly reduced IFNγ and TNF production compared to controls (Figures 2J and S3D), suggesting that prior respiratory viral infections diminished anti-tumor T cell responses. Taken together, these data suggest that prior viral pneumonia reprograms the local lung environment into a pro-tumor state, characterized by an increased presence of pro-tumor myeloid cells and a decreased anti-tumor T cell immunity.

Recent evidence has revealed important roles of tumor-infiltrating neutrophils in promoting lung cancer growth.^18^ To gain deeper insights into neutrophil heterogeneity and function in this context, we subclustered neutrophils from our dataset and identified 8 clusters (Figures 2K and S2L). Among them, clusters 0, 2, 3, 4, and 7 were present in non-tumor bearing lungs. Conversely, clusters 1, 5 and 6 were highly enriched in tumor-bearing lungs and were thus designated as TANs. Notably, cluster 1 was the most abundant TAN subset in IAV tumor-bearing lungs at both early and late stages (Figure 2L). It was transcriptionally defined by elevated expression of pro-tumorigenic genes such as *Il1a*, *Ccl3*, *Vegfa* and *Cd63* (Figures S2L, and S2M). Cluster 6, which expressed *Cxcl3*, *Xbp1* and *Tnfrsf23*, was also preferentially present in IAV tumors (Figures S2L and S2M). Interestingly, cluster 1 also exhibited high expression of *Siglecf* (Figures 2M and S2N), a marker previously associated with pro-tumor neutrophils.^39–41^ Flow cytometry analysis showed that SiglecF^hi^ neutrophils predominantly resided in the lung parenchyma, with minimal presence in the blood, indicating that they may be developed locally and interact with the tumor (Figures 2N-2P, and S3E-S3G). Consistent with this notion, immunofluorescence staining for SiglecF^+^ MPO^+^ neutrophils confirmed a marked increase in tumor-bearing lungs of mice previously infected with IAV or SARS-CoV-2 compared to their respective PBS-treated controls (Figures 2Q-2S and S2O).

Previous studies have demonstrated that SiglecF^hi^ neutrophils exhibit pro-tumor functions in lung cancer.^39–41^ Consistent with this, differential gene expression analysis revealed that *Siglecf^hi^*neutrophils (c1) upregulated genes associated with myeloid cell recruitment, tumor proliferation and immunosuppression compared to those of *Siglecf^low^*neutrophils (Figure S2P). Pathway enrichment further showed that *Siglecf^hi^*neutrophils were enriched pathways related to glycolysis, hypoxia, TNF, TGF-β, and mTORC signaling, as well as Myc and E2F targets and programs negative regulation of cell death. In contrast, *Siglecf^low^* neutrophils were linked to neutrophil activation, T and NK cell activation, defense responses and IFN signaling (Figure 2T). These differences were recapitulated in comparisons between IAV- and PBS-derived *Siglecf^hi^*TANs (Figures S2Q-S2S), supporting the notion that prior viral infection reprograms neutrophils toward a pro-tumor phenotype. To functionally test if *Siglecf^hi^*TANs from prior infected mice possess increased tumor-promoting capacity, we adopted a tumor and neutrophil co-transfer model that were previous reported.^21^ KP tumor cells were subcutaneously (s.c.) implanted and co-injected with sorted SiglecF^hi^ or SiglecF^low^ neutrophils isolated from IAV_KPi.t. or PBS_KPi.t. lungs. Mice received intratumoral neutrophils boosts on days 3 and 7 post-implantation (Figure 2U). Tumors co-injected with IAV-derived SiglecF^hi^ neutrophils grew significantly faster than those co-injected with PBS-derived SiglecF^hi^ neutrophils or either SiglecF^low^ controls and had the greatest tumor mass at the end point (Figure 2U).

A recent study in pancreatic ductal adenocarcinoma (PDAC) described a deterministic neutrophil reprogramming into a terminal T3 state, localizing to the hypoxic-glycolytic niches where they promote angiogenesis and support tumor growth.^21^ Applying the T3 gene signature to our neutrophil clusters, we found that the overall T3 signature was mostly enriched in Neu_1 *Siglecf*-expressing (Figure 2V; Table S2).

Moreover, cell-cell communication analysis (*CellChat)* further revealed that the Neu_1 *Siglecf-*expressing population exhibited the strongest predicted interactions with the progenitor and tumor cell subsets (KAC, early and later tumor cells) (Figures 2W and S2T). These results suggest that in the lung cancer setting, *Siglecf^hi^* neutrophils closely resemble the T3 function state and are key mediator of tumor-promoting activity. To explore the clinical relevance of this findings, we derived a *Siglecf^hi^* neutrophil gene signature based on the top 100 differentially expressed genes (DEGs) between IAV- and PBS-derived *Siglecf^hi^* TANs (Table S2). Applying this signature to LUAD patient data from TCGA, we found that high signature expression of *Siglecf^hi^* neutrophils from prior IAV infected mice correlated with significantly worse overall survival (Figure 2X). This result held even when using a more conservative gene set composed of shared *Siglecf^hi^* markers between mouse (this study) and human neutrophils^24^ (Figure 2X).

Collectively, these findings indicate that prior viral infections program subsequent pro-tumor lung microenvironment characterized by the high presence of tumor promoting SiglecF^hi^ TANs.

### Prior infections epigenetically reprogram lung cells to favor tumor growth

Neutrophils are short-lived immune cells and often known their ability to undergo progenitor reprogramming in the bone marrow (BM) following exposure to various insults.^42,43^ To determine whether the protumor effects of neutrophil in virus-infected mice were due to long-term reprogramming of BM neutrophil progenitors, we conducted a BM transfer experiment. BM cells from IAV- or PBS-infected donor mice (21 d.p.i.) were transplanted into lethally-irradiated naïve recipients, followed by KP tumor cell i.t. inoculation. Notably, there was no significant difference in tumor growth observed between mice transferred from prior-infected or non-infected mice (Figures 3A and 3B), indicating that BM neutrophil progenitors from prior infected mice did not enhance tumor growth. Consistently, s.c. implantation of KP cells into prior IAV-infected mice also did not result in enhanced tumor growth (Figures 3C, S4A and S4B). These findings suggest that the pro-tumor effects observed in our models are likely lung-specific rather than stemming from systemic BM reprogramming.

**Figure 3.**
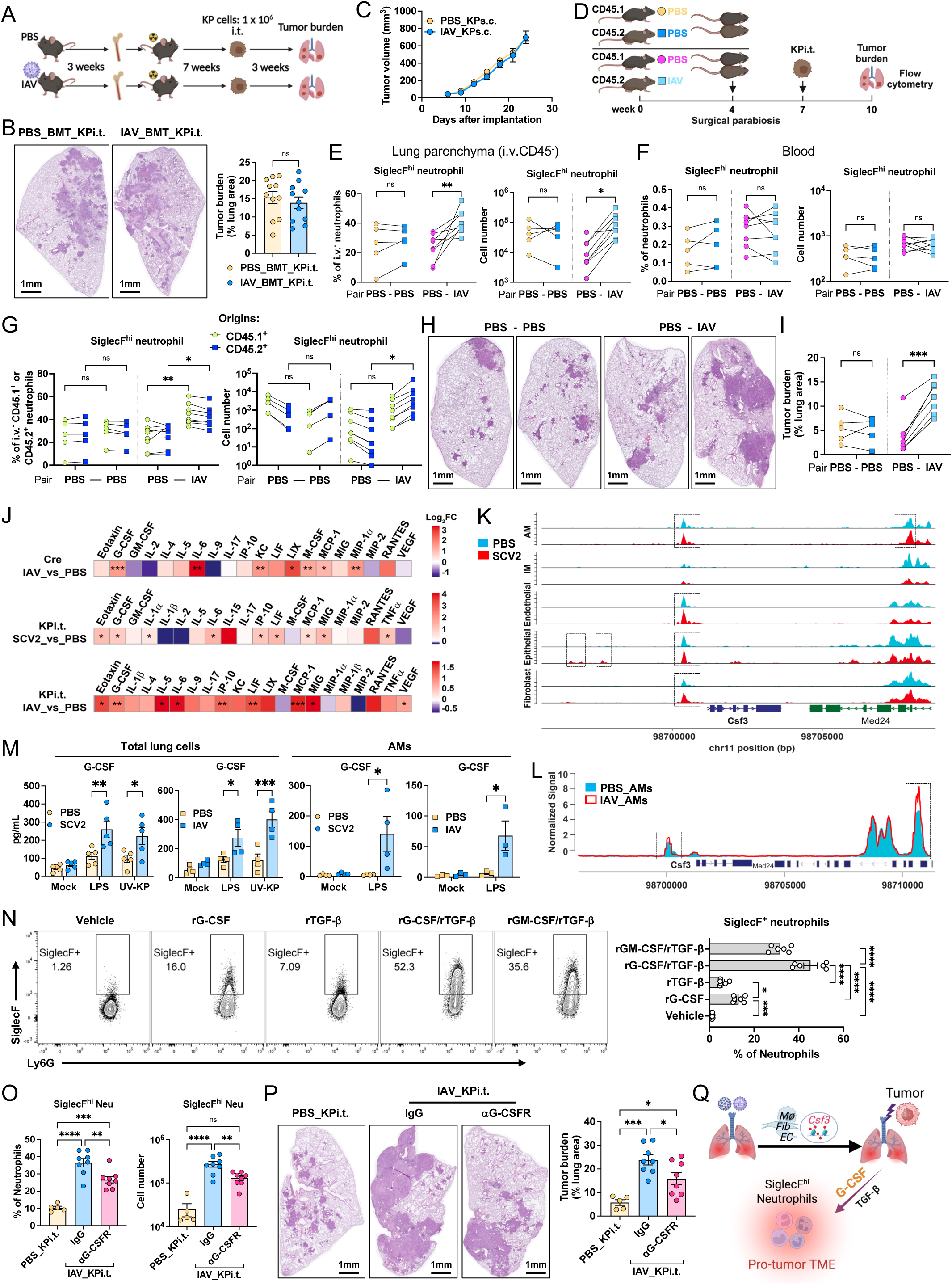
Epigenetically imprinted lung microenvironment by viral infection promotes lung tumor growth. (A) Schematic of experimental setup for bone marrow transplantation and tumor implantation. (B) Representative image of tumor-bearing lung lobes (left) and quantification of tumor burden (right) at week 3 in PBS- or IAV-bone marrow transferred KPi.t. mice. (C) Tumor growth curves of PBS- or IAV-infected mice s.c. implanted with KP tumor cells. (D) Schematic of the tumor parabiosis experiment. Wild-type female CD45.1^+^ and CD45.2^+^ congenic mice were treated with PBS or infected with IAV, respectively, and maintained for 4 weeks prior to surgical parabiosis (forming PBS-IAV pairs). PBS-PBS pairs served as controls. After 3 weeks of parabiosis recovery, both mice in each pair were i.t. inoculated with KP tumor cells. Tumor burden and neutrophil populations were analyzed 3 weeks later. (**E and F**) Frequency and cell number of SiglecF^hi^ neutrophils in the lung parenchyma (E) and blood (F) among parabiosis pairs. (**G**) Frequency and cell number of SiglecF^hi^ neutrophils in the lung parenchyma of parabiosis pairs, stratified by CD45.1^+^ and CD45.2^+^ origin to distinguish host- vs partner-derived cells. (**H and I**) Representative image of tumor-bearing lung lobes (H) and quantification of tumor burden (I) at week 3 among parabiosis pairs. (J) Heatmap showing upregulated cytokines and chemokines in BAL from IAV_Cre, IAV_KPi.t. and SCV2_KPi.t. mice compared to their respective PBS-treated controls. (K) Representative scATAC-Seq genomic browser tracks showing chromatin accessibility at the *Csf3* locus in AM, IM, epithelial cells, fibroblasts and endothelial cells from PBS- and SCV2-infected lungs at 28 d.p.i. (L) Representative bulk ATAC-Seq genomic browser tracks displaying chromatin accessibility at the *Csf3* locus in AM isolated from BAL of PBS- and IAV-infected mice at 35 d.p.i. (M) G-CSF production by SCV2- or IAV-infected lung cells or AMs at 35 d.p.i. following overnight stimulation with LPS or UV-irradiated KP cells. Culture supernatants were collected and analyzed by ELISA. (N) Flow cytometry dot plots showing the induction of SiglecF^+^ neutrophils following recombinant G-CSF or TGF-β treatment, as well as G-CSF/TGF-β and GM-CSF/TGF-β co-treatment (left), and the proportion of SiglecF^+^ neutrophils (right). (O) Blockade of G-CSF receptor (G-CSFR) reduced both the frequency and number of SiglecF^hi^ neutrophils in tumor-bearing lungs of mice previously infected with IAV. (P) Representative image of tumor-bearing lung lobes and quantification of tumor burden from (O). (Q) Schematic model illustrating how prior severe respiratory viral infection epigenetically reprograms lung microenvironment via G-CSF production, promoting the local accumulation of SiglecF^hi^ neutrophils and contributing to accelerated tumor growth. Data represent two independent experiments or pooled from two experiments. Graphs display mean ± SEM. Statistical significance was assessed by Mann-Whitney test (B and J), two-way ANOVA (E, F, G, and I), and one-way ANOVA (N). (ns *p* > 0.05, \**p* < 0.05, \*\**p* < 0.01, \*\*\**p* < 0.001, \*\*\*\**p* < 0.0001)

To directly test this hypothesis, we conducted a parabiosis experiment to surgically join the circulation of non-infected (CD45.1^+^) mice with prior-infected (CD45.2^+^) congenic mice (Figure 3D). Following three weeks of parabiosis, chimerism of circulating immune cells, including T and B cells, was achieved (Figure S4C). Both parabionts were then inoculated i.t. with KP tumor cells. Strikingly, IAV-tumor-bearing mice from PBS-IAV pairs harbored significantly more total neutrophils and SiglecF^hi^ neutrophils in the lung parenchyma than their PBS-tumor-bearing mice (Figures 3E and S4D), whereas blood neutrophil levels remained comparable (Figures 3F and S4E).

Importantly, the elevated SiglecF^hi^ neutrophils were independent of their origins, as both CD45.2^+^ mouse- and CD45.1^+^ mouse-derived neutrophils showed similar expansion in IAV-tumor bearing mice (Figure 3G), reinforcing the dominant role of the virus-conditioned lung microenvironment. PBS-PBS pairs showed no significant differences in any of these parameters. Furthermore, IAV parabionts of PBS-IAV pairs exhibited significantly higher tumor burden than PBS parabionts, while PBS-PBS pairs showed comparable tumor growth among the paired mice (Figures 3H and 3I). These data collectively support the conclusion that prior infection creates a lung niche that drives the local accumulation and reprogramming of pro-tumor neutrophils.

To understand how the infected lung environment supports enhanced neutrophil recruitment and reprogramming, we profiled cytokines and chemokines in BAL from tumor bearing mice with or without prior infection. Across both genetic and orthotopic transfer tumor models, we found significant upregulation of G-CSF, IL-6 and certain chemokines such as MCP-1 (Figure 3J), which have been shown to promote the development, survival, recruitment and/or differentiation of neutrophils.^18,20^ These data suggest that prior infection may epigenetically program heightened cytokine production to facilitate local neutrophil recruitment and pro-tumor TAN differentiation. To confirm this idea, we performed scATAC-seq on total lung cells from SARS-CoV-2-infected (28 d.p.i.) and PBS-treated mice. Unsupervised clustering of chromatin accessibility profiles identified major immune and stromal populations (Figure S4F). At the epigenetic level, we found mildly to moderately increased chromatin accessibility at the *Csf3* locus in macrophages, epithelial cells, fibroblasts and endothelial cells from SARS-CoV-2 infected lungs (Figure 3K). Similar increases were observed at the *Il6*, *Il1b*, *Cxcl1* and *Cxcl5* locus across several cell types (Figure S4G). To identify regulators driving this epigenetic remodeling, we performed motif enrichment and regulon activity analysis.

NF-κB and AP-1 motifs were enriched and transcriptionally active across several cell populations, indicating persistent inflammatory signaling. STAT3 and STAT4 activity was also elevated in multiple compartments, consistent with sustained cytokine signaling downstream of IL-6 and G-CSF (Figure S4I).

Previous studies have shown that influenza infection was associated with the “training” of AMs for their enhanced sensitivity to a secondary stimuli.^44,45^ To this end, we also performed bulk ATAC-seq on AMs isolated from IAV-infected mice and PBS controls. AMs from infected mice exhibited increased chromatin accessibility at the *Csf3* locus compared to AMs from uninfected mice (Figure 3L). These results were consistent with previously published ATAC-seq data from IAV-infected AMs,^44^ which showed enhanced *Csf3* locus accessibility in both resident and monocyte-derived AMs (Figure S4H). Together, these data reveal that prior respiratory viral infection induces long-lasting epigenetic remodeling of lung immune and structural cells, programming inflammatory cytokine networks that promote tumor growth and immune suppression.

To test whether this epigenetic priming may translate into increased cytokine production following secondary exposure, we performed *in vitro* restimulation assay. Total lung cells or AMs isolated from SARS-CoV-2- or IAV-infected (35 d.p.i.) and their respective PBS-treated mice were restimulated with LPS or UV-irradiated KP cells. Viral infection-experienced lung cells and AMs produced significantly more G-CSF in response to LPS or irradiated KP cell stimulation, whereas basal G-CSF levels in unstimulated (vehicle) cells were similar between groups (Figure 3M). These data indicate that prior infection induces a lasting epigenetic state in lung stromal and immune cells, enhancing their ability to produce neutrophil-promoting cytokines upon tumor challenge.

Given this, we next reasoned how infection-induced signals promote the emergence of SiglecF^hi^ neutrophils. Previous studies have shown that SiglecF expression on neutrophils can be induced by GM-CSF and TGF-β in renal fibrosis or cancer settings.^46,47^ However, since GM-CSF or TGF-β was not upregulated with prior infection (Figures 3J and S4J), we tested whether G-CSF could facilitate SiglecF expression on neutrophils. G-CSF alone was able to induce notable SiglecF expression, and in combination with TGF-β elicited strongest induction, exceeding that of GM-CSF+TGF-β (Figures 3N and S4K). Consistently, intranasally administration of G-CSF plus TGF-β into naïve mice enhanced both total neutrophil recruitment and SiglecF expression in the lung (Figures S4L and S4M). To test if blocking local G-CSF signaling could counteract infection-driven neutrophil reprogramming and tumor progression, we treated IAV-infected mice with anti-G-CSFR antibody or IgG isotype control intranasally starting one day prior to KP tumor cell inoculation. Local anti-G-CSFR treatment significantly reduced SiglecF^hi^ neutrophils in the lung (Figure 3O) and resulted in a decrease in tumor burden compared to mice treated with IgG antibody (Figure 3P). Altogether, our findings indicate that prior respiratory viral infections induce long-lasting, epigenetically encoded changes in the lung microenvironment that promote cytokine-mediated neutrophil recruitment and reprogramming. This lung-specific imprinting by prior viral infection supports the expansion of SiglecF^hi^ neutrophils and creates a permissive local environment for tumor growth (Figure 3Q).

### Therapeutic strategies targeting pro-tumor environment

Next, we sought to examine whether specifically targeting neutrophils could ameliorate lung tumor burden after prior viral infection. To this end, we evaluated two complementary strategies aimed at inhibiting neutrophil recruitment and function: 1) neutrophil depletion via anti-Ly6G antibody and 2) blockade of the CXCR2 signaling axis using the small molecule inbibitor Reparixin (REP). We treated virus-infected mice with either anti-Ly6G antibody or REP one day prior to KP tumor cell i.t. inoculation and continued every other day until the experiment endpoint, IgG isotype antibody or DMSO vehicle was applied as respective controls (Figure 4A). Both interventions led to partial, yet significant, reduction of lung neutrophils as confirmed by flow cytometry (Figures S5A and S5B). Importantly, anti-Ly6G and REP treatments resulted in significantly reduced tumor burden in SARS-CoV-2-infected mice (Figures 4B-4E) and improved survival in tumor-bearing IAV-infected mice (Figure 4F). These findings demonstrate that the interference with neutrophil responses could limit tumor progression in post-infection settings.

**Figure 4.**
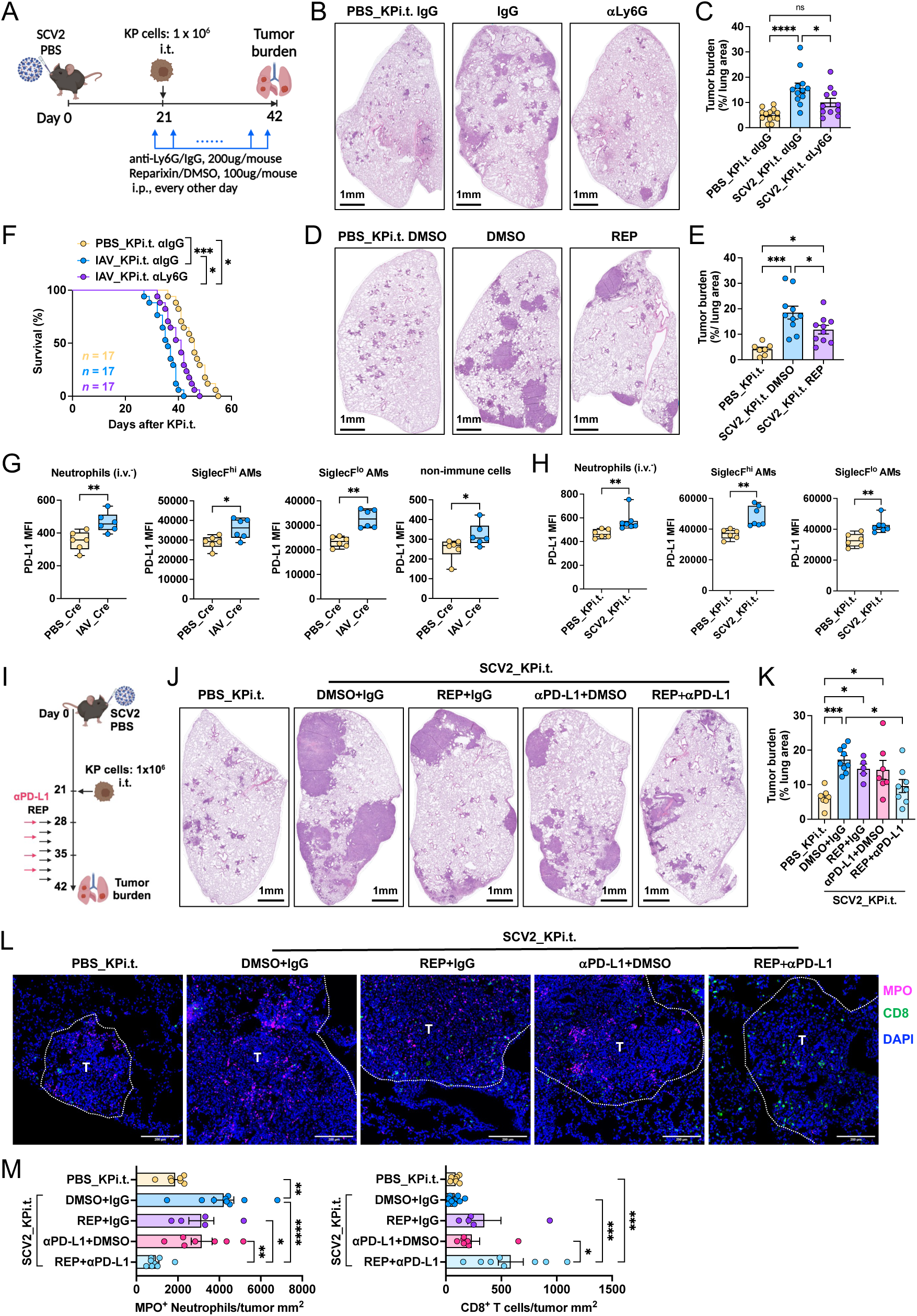
Therapeutic targeting pro-tumor neutrophils and PD-L1 ameliorates lung tumor growth after infection. (A) Experimental design of neutrophil depletion strategy. Mice were infected with SCV2 for 21 days before inoculated with KP tumor cells. Anti-Ly6G antibody or CXCR2 inhibitor Reparixin (REP) was administered starting one day prior to tumor implantation and continued every other day until the experiment endpoint. (**B and C**) Representative image (B) and quantification of tumor burden (C) in lungs of PBS- or SCV2-infected mice treated with anti-Ly6G. (**D and E**) Representative image (D) and tumor burden quantification (E) in lungs of PBS- or SCV2-infected mice treated with Reparixin (REP). (F) Survival curve of control and anti-Ly6G-treated tumor-bearing prior IAV-infected mice. (**G and H**) MFI of PD-L1 expression on indicated cells from tumor-bearing lungs of IAV_Cre (G) or SCV2_KPi.t. (H) mice compared to their respective PBS-treated controls. (**I**) Experimental design for neutrophil depletion combined with anti-PD-L1 immunotherapy in the SCV2_KPi.t. model. (**J and K**) Representative image (J) and quantification of tumor burden (K) in SCV2-infected mice treated with REP alone, anti-PD-L1 alone, or in combination. (**L**) Representative immunofluorescence images showing MPO and CD8 staining in tumor sections from (J). White dished lines outline tumor regions. Scale bar, 200 μm. (**M**) Quantification of MPO^+^ neutrophils and CD8^+^ T cells from (L). Data represented or pooled from two (E, G, H, K, and M) and three (C and F) independent experiments. Graphs display mean ± SEM. Statistical significance was assessed by Mann-Whitney test (G and H), one-way ANOVA (C, E, K and M), and Log-rank Mantel-Cox test (F). (ns *p* > 0.05, \**p* < 0.05, \*\**p* < 0.01, \*\*\**p* < 0.001, \*\*\*\**p* < 0.0001)

We reasoned that viral infection not only amplifies neutrophilic inflammation but also promotes immune evasion through checkpoint upregulation. PD-L1 (CD274), a key immunosuppressive molecule, is known to be expressed on tumor cells and various suppressive immune subsets, including TANs and TAMs.^26,48^ Our scRNA-seq data revealed elevated *Cd274* expression in multiple immune cell types, including neutrophils, AMs, IMs, monocytes, and dendritic cells in IAV_T12 tumors compared to PBS_T12 tumors (Figure S5C). This upregulation was validated by flow cytometry in both SARS-CoV-2 and IAV infection tumor models, showing significantly increased PD-L1 protein levels in AMs, neutrophils, and non-immune cells in virus-infected lungs (Figures 4G, 4H, S5D and S5E). These data suggest that post-viral TMEs are not only neutrophil-rich but also likely exhibited increased PD-L1-mediated T cell inhibition, consistent with previous CD8 T cell dysfunction data (Figures 2I and 2J). Given the concurrent elevation of neutrophils and PD-L1-driven immune suppression, we next tested whether combined neutrophil inhibition and PD-L1 blockade could be used therapeutically to diminish lung cancer growth after viral pneumonia. To this end, prior SARS-CoV-2 infected tumor bearing mice were treated with REP alone, anti-PD-L1 antibody alone, or a combination of both, starting one week after tumor inoculation (Figure 4I). While each monotherapy mildly reduced tumor burden in this therapeutic time window (1 week after tumor inoculation), the combination of CXCR2 inhibition and PD-L1 blockade significantly reduced tumor burden (Figures 4J and 4K). This indicates that simultaneous blockade of neutrophil-driven immunosuppression and T cell checkpoint inhibition is required to therapeutically dampen the tumor-promoting effects established in virus-primed lung microenvironment. Mechanistically, the dual treatment diminished the pro-tumor TME and enhanced antitumor T cell immunity, as evidenced by a significant decrease in MPO^+^ neutrophil accumulation and a notable increase in CD8^+^ T cell infiltration within the tumor-bearing lungs (Figures 4L and 4M).

## Discussion

Respiratory viral infections are among the most prevalent insults to the lung, yet their long-term consequences on cancer susceptibility remain underexplored. Here, we demonstrate that prior severe respiratory viral infections, including SARS-CoV-2 and IAV, can immunologically and epigenetically reprogram the lung microenvironment to promote tumor development and progression. Through a combination of retrospective human data, diverse murine lung cancer models and single cell multi-omics analysis, our study establishes a mechanistic link between prior severe viral infection and heightened lung cancer risk.

Our retrospective cohort analysis revealed a significant association between prior hospitalization for COVID-19 and increased lung cancer incidence, independent of smoking history. Interestingly, this risk was not observed and was slightly reduced in patients with mild/moderate infections (Figure 1B). This divergence likely reflects the distinct immunological consequences of infection severity. Severe respiratory viral infections, including SARS-CoV-2 and IAV, are known to induce persistent epithelial injury, unresolved inflammation and fibrotic remodeling in the lung, features that persist years after viral clearance.^27,29,31,49–51^ These tissue abnormalities likely create a microenvironment conducive to malignant transformation through chronic injury, dysregulated repair, and immune dysfunction, all of them are hallmarks of early tumorigenesis.^52^ Supporting this, our findings converge with a prior report linking cumulative influenza exposure to elevated lung cancer risk,^8^ and with a recent study demonstrated that influenza and SARS-CoV-2 infections reactivate dormant metastatic cells in the lung.^53^ Further reinforcing this concept, a recent large-scale epidemiological study showed that exposure to PM2.5 air pollutants was associated with significantly increased incidence of EGFR-mutant LUAD (HR=1.08), particularly in non-smokers.^54^ This tumor-promoting effect was shown to be inflammation-driven, involving reprogrammed AT2 cells into a progenitor-like cell state that fuels tumorigenesis.

Together, these data suggest that diverse inflammation insults, including viral pneumonia and environmental pollutants, can act as a priming event that influence cancer risk months or even years after apparent clinical recovery. These mechanisms may extend beyond lung cancer to metastasis-prone malignancies such as breast cancer, raising the possibility that PASC-associated immune remodeling could facilitate metastatic outgrowth, a question warrants future investigation.

In this context, the post-COVID-19 era, with tens of millions of people globally experiencing long-term pulmonary sequelae,^1^ represents a pivotal moment for rethinking cancer risk in the setting of infection-induced tissue programming. These findings carry significant implications for clinical care. Individuals recovering from severe viral pneumonia, particularly those with smoking history, may benefit from enhanced lung cancer surveillance. In addition, preventing severe infection through vaccination may confer indirect cancer protection benefits. This concept is supported by our experimental data, where mRNA-based SARS-CoV-2 vaccination mitigated the tumor-promoting effects of prior infection.

One of the most striking observations in our study was the emergence of more advanced tumor states in the LUAD GEMM mice previously exposed to viral infection. Notably, the IAV-infected group exhibited an increased abundance of KRT8^hi^ alveolar intermediate cells (KACs) and LUAD clusters, suggesting that prior epithelial injury fundamentally alters the trajectory of epithelial cell differentiation and tumorigenesis. This concept aligns with accumulating evidence that AT2 cells can be hijacked by oncogenic signals in the context of tissue repair, in particular, sustained NF-κB activation in mutant AT2 cells drives a regenerative-like program that delays terminal differentiation and facilitates transformation.^55^ Our pseudotime and CytoTRACE analyses similarly showed that KACs lie at the bifurcation between AT1 and LUAD fates, echoing a recent single-cell atlas of early LUAD, where KRT8^+^ intermediate cells act as a critical transitional population in the AT2-to-tumor trajectory.^35^ This suggests that epithelial cells activated by prior injury remain in a primed, plastic state that is permissive to tumorigenesis when exposed to oncogenic stimuli, although the precise mechanisms and causal links warrant further in-depth investigation.

Our bone marrow chimera and parabiosis experiments indicate that immune (neutrophils) programming in the post-viral lung is lung-specific rather than hematopoietic, aligning with recent findings that local tissue environments can educate neutrophils independently of systemic signals.^19,46,56^ Within this locally remodeled immune landscape, we identified a functionally distinct subset of TANs, marked by high SiglecF expression, that are expanded in the virus-experienced tumor-bearing lungs.

Consistent with previous findings,^39^ SiglecF^hi^ TANs exhibit a transcriptional program enriched for pro-tumor pathways, features that also mirror the T3 terminal neutrophil state described in pancreatic cancer and other solid tumors.^21^ While SiglecF^hi^ neutrophils have been linked to tumor-induced systemic remodeling of granulopoiesis, including osteoblast activation and partial transcriptional priming in circulation,^39^ our findings suggest that local infection-primed cues in the lung further instruct this tumor-promoting phenotype. Of note, their accumulation was associated with reduced CD8⁺ T cell infiltration and diminished effector gene expression in the tumor microenvironment, suggesting a potential role of them in shaping local immune suppression. However, the mechanisms by which SiglecF^hi^ TANs influence T cell function remain unclear currently. Possible mediators include PD-L1 expression, production of arginase or reactive oxygen species, or metabolic competition, as implicated in other neutrophil-driven tumor contexts.^18,20^ Future work with functional perturbation studies will be essential to dissect whether these cells act through direct contact or paracrine signaling.

Importantly, SiglecF^hi^ TANs are unlikely to act alone. Tumor-associated macrophages (TAMs) also represent a dominant immunosuppressive population in lung cancer,^57^ and in our study, late-stage IAV-experienced tumors exhibited increased enrichment of a pro-tumor TAM signature. This finding aligns with extensive literature showing that TAMs contribute to immune evasion, therapeutic resistance, and angiogenesis in LUAD.^58–60^ While TANs and TAMs may share overlapping immunosuppressive functions, they are recruited and shaped by distinct cytokine and chemokine cues and may act in parallel to sustain an immunosuppressive tumor microenvironment.^57^ Whether therapeutic targeting of one myeloid population results in compensatory remodeling or expansion of the other remains further investigation.

Together, our findings indicate that epigenetic reprogramming of the lung environment potently drives the heightened tumor growth observed after viral pneumonia. We also identify a neutrophil-centered therapeutic approach that can counteract immunosuppression and curb the surge in tumor development following infection. Beyond these mechanistic and therapeutic insights, our results underscore the importance of vigilant cancer screening and prevention strategies for COVID-19 survivors, especially those with persistent lung sequelae or additional risk factors such as smoking or pollutant exposure.

## RESOURCE AVAILABILITY

### Lead contact

Jie Sun

### Materials availability

Materials used in the manuscript are available upon reasonable request to the Lead author.

### Data and code availability

Data and code will be deposited and available upon the publication of the manuscript.

## ACKNOWLEDGMENTS

We thank Dr. Nikil Josh for the KP cell lines. Schematic diagram in the manuscript were created with BioRender.com. Data for this manuscript were generated using the Flow Cytometry Research Histology Core Facilities, Biorepository and Tissue Research Facility and Biomolecular Analysis Facility at the University of Virginia. The study was in part supported by the US National Institutes of Health(NIH) grants AI147394, AG069264, AI112844, HL170961, AI176171 and AG090337 to J.S., UVA Pinn Scholar to J.S., UVA Comprehensive Cancer Center (CCC) Collaborative Grant to J.S. and K.P., U01CA224293 and UVA Shannon Fellowship to K.P., UVACCC Lung TRT Pilot Grant to J.S., NIH F31HL170746 and T32AI007496 to H.N., NIH

T32CA009109 to S.P.Y., American Lung Association Catalyst Grant to I.S.C., T32GM139787-01 and UVA Parsons-Weber-Parsons Fellowship to N.A.K.

## AUTHOR CONTRIBUTIONS

J.S. supervised the project. W.Q. & J.S. conceived the overall project. W.Q., X.W., J.S. designed the experimental strategy and analyzed data. W.Q., X.W., A.J.B., X.Y., J.Y., S.P.Y, E.V.Y., Y.P., C.L., G.A.S, J.T., H.N., P.C., J.M.S., K.P., W.C., I.S.C. performed experiments, analyzed data, or contributed critical reagents to the study. W.Q. & J.S. wrote the original draft. All authors read, edited, and approved the final manuscript.

## STAR★METHODS

### KEY RESOURCES TABLE

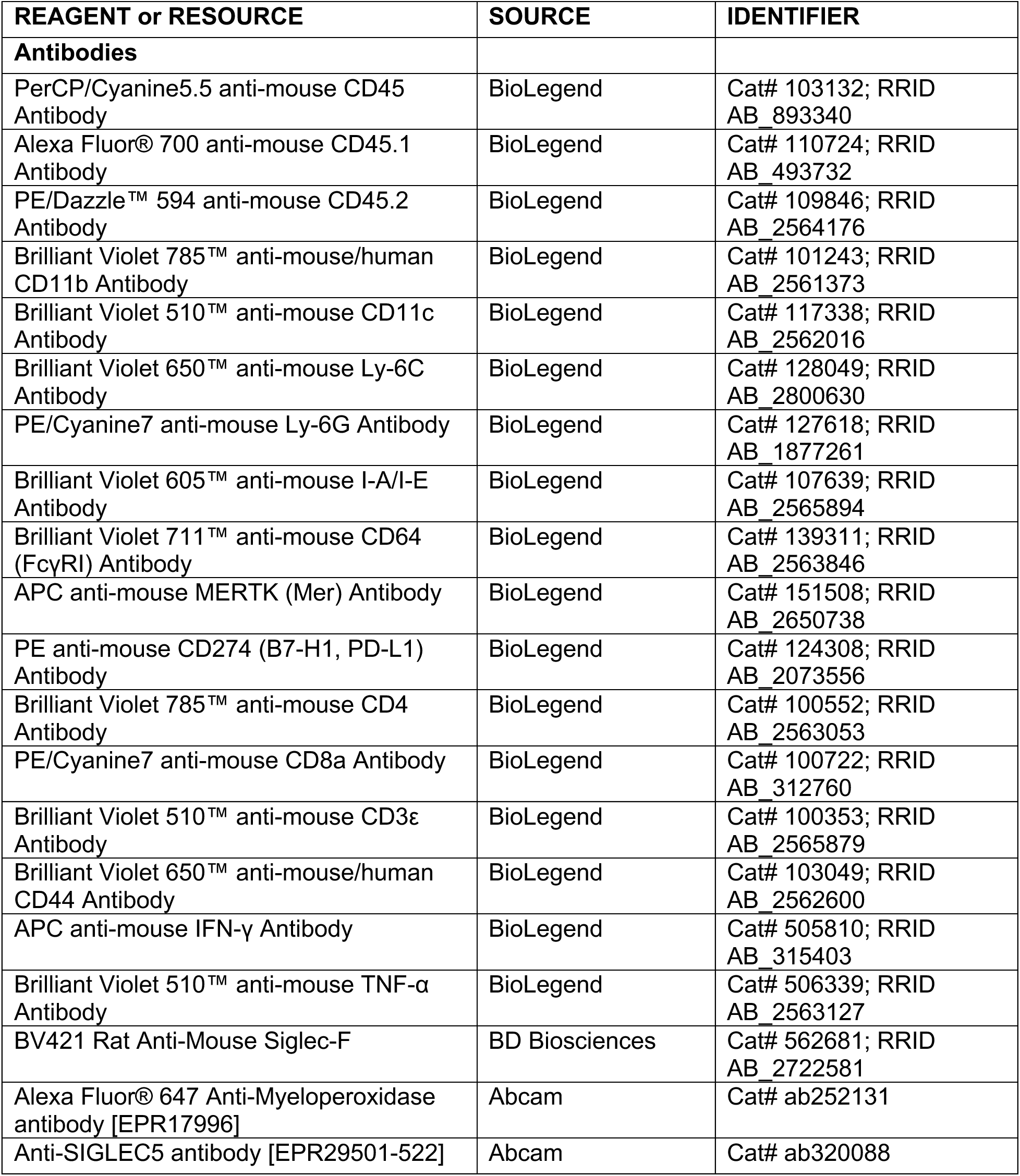
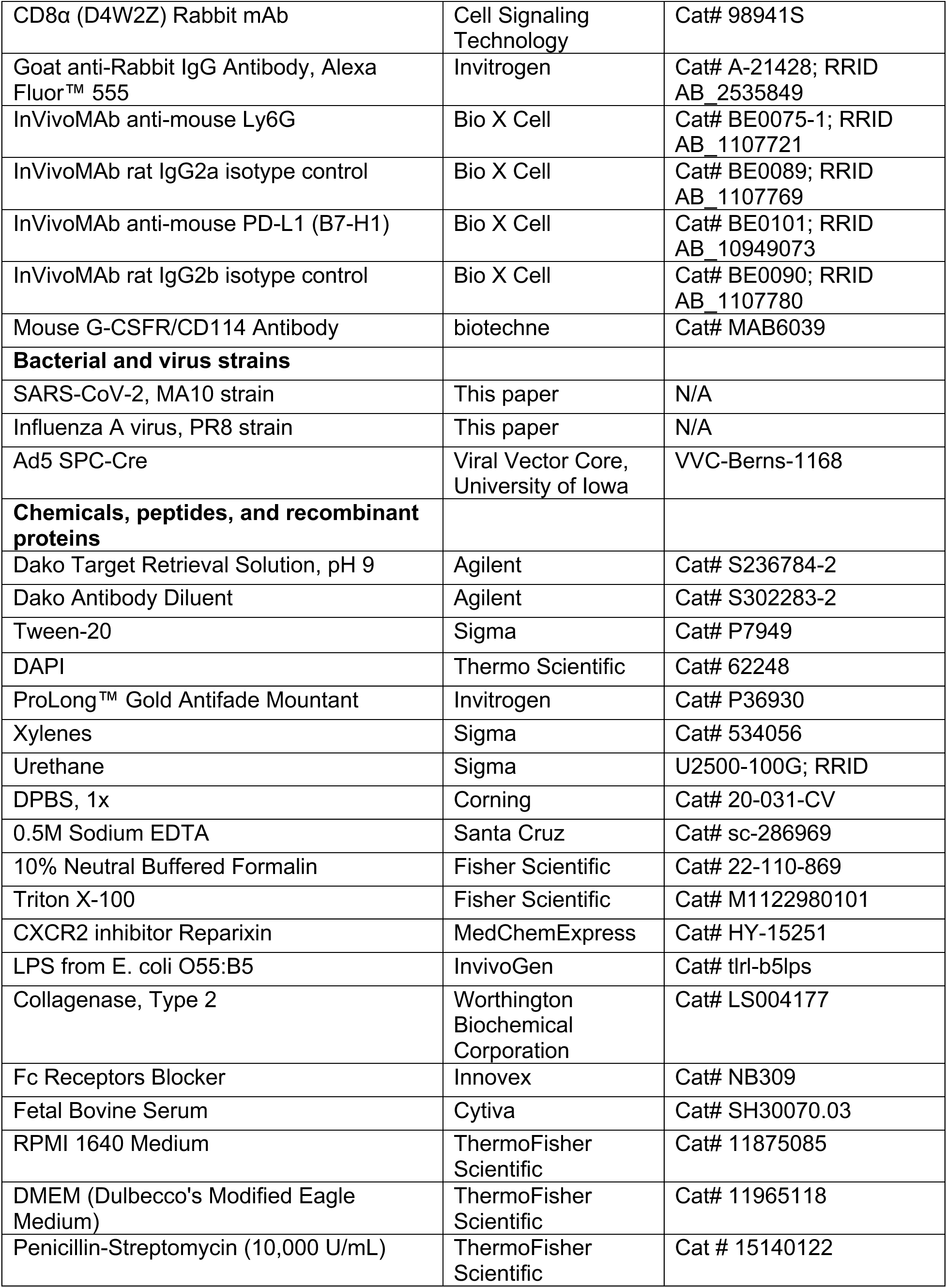
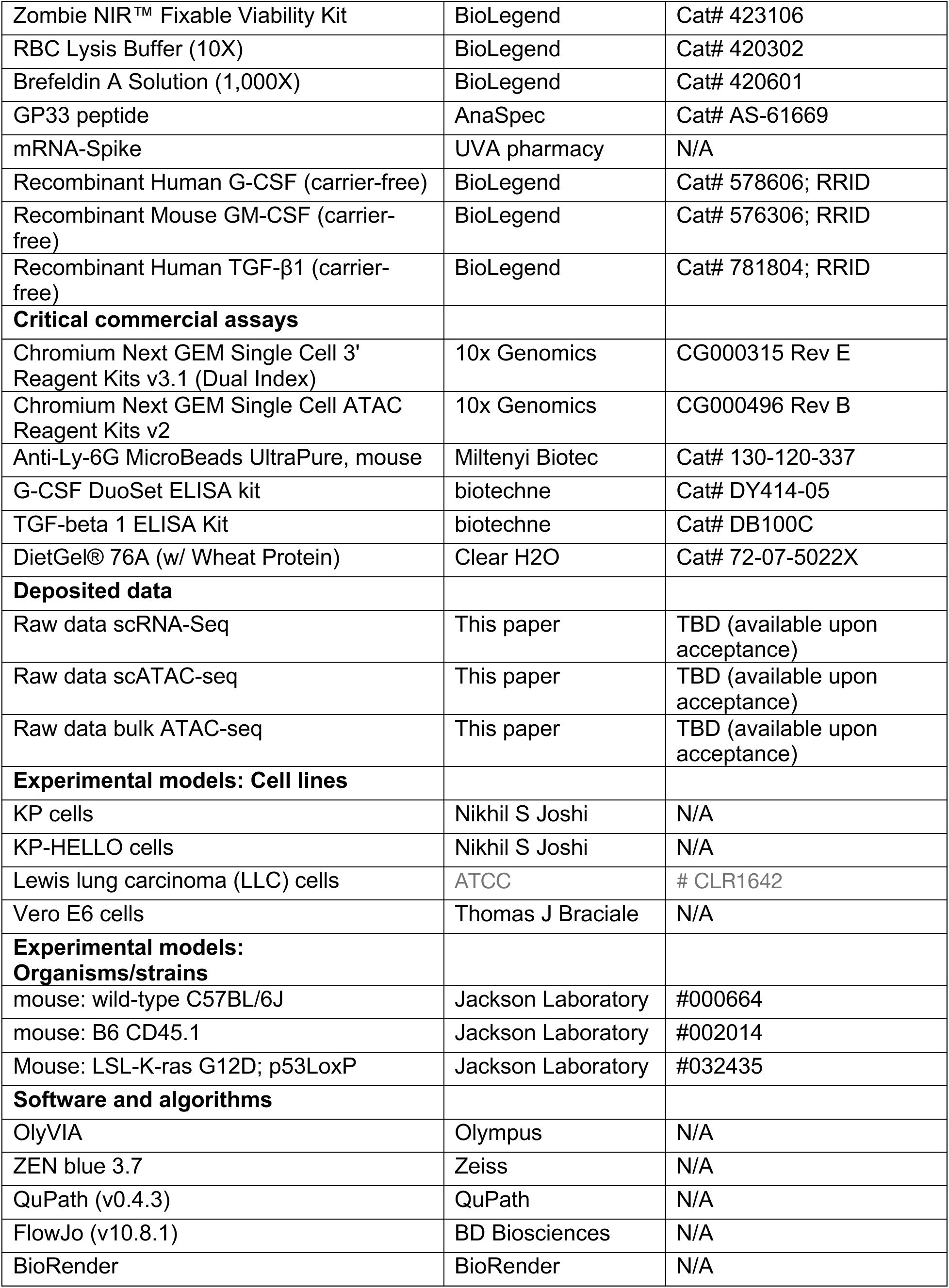
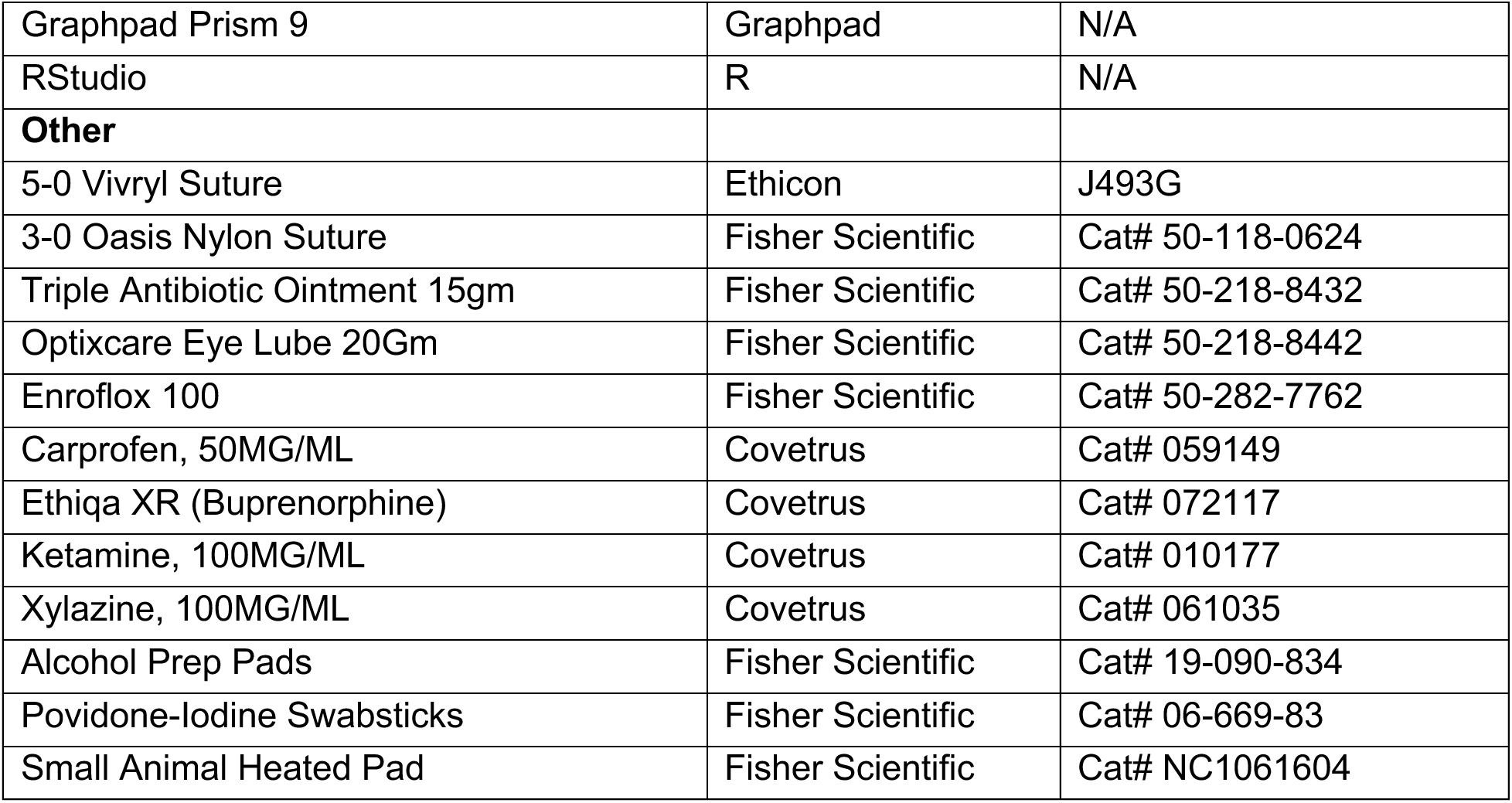

### EXPERIMENTAL MODEL AND STUDY PARTICIPANT DETAILS

#### Human epidemiological studies

We conducted a large-scale, retrospective cohort study using data from Epic Cosmos, a dataset created in collaboration with a community of Epic health systems representing more than 296 million patient records from over 1704 hospitals and 39,900 clinics from all 50 states, D.C., Lebanon, and Saudi Arabia. Data were harmonized and de-identified across participating health systems and accessed in accordance with institutional data-sharing agreements. The study cohort included 44,229,908 adults aged ≥45 years who were alive and without a known history of cancer as of January 1, 2022. The primary exposure was a recorded diagnosis of SARS-CoV-2 infection prior to January 1, 2022. Infection severity was stratified into ambulatory (mild/moderate illness) and hospitalized (severe illness) based on the encounter type of the index COVID-19 diagnosis. The primary outcome was incident cancer, defined by a new diagnosis code accompanied by staging documentation within the EHR system.

We estimated hazard ratios (HRs) and 95% confidence intervals (CIs) using multivariable Cox proportional hazards models, adjusting for age at cohort entry, sex, and smoking history. Analyses were performed using R version 4.1.0. Patients were followed from January 1, 2022 until the earliest of cancer diagnosis, death, loss to follow-up, or end of the study period.

#### Mice

Wild-type C57BL/6J female and male mice (10-week-old) were originally purchased from Jackson Laboratory and housed and inbred in our animal facility. Wild-type C57BL/6J male mice (25-week-old) were purchased from Jackson Laboratory and were housed for at least two weeks before used for infection. Transgenic mouse model of lung adenocarcinoma Kras^LSL-G12D/+^ Trp53^flox/flox^ mice (referred to as KP mice) were purchased from Jackson Laboratory and housed and inbred in our animal facility.

Among them, 12- to 15-week-old female and male KP mice were used for IAV infection, while 6-mo-old male mice were used for SARS-CoV-2 MA10 infection. All animal experiments were performed in animal facilities at the University of Virginia (UVA). The animal experiments were approved by UVA Institutional Animal Care and Use Committee. All work with SARS-CoV-2 infections were approved under Animal Biosafety Level 3 (ABSL3) conditions and were performed with approved standard operating procedures and safety conditions by the UVA Institutional Review Board.

#### Cell lines and virus

KP cells derived from tumor-bearing lungs of KP mice, as well as KP cells engineered to express the model antigen HELLO for B and T cell recognition (referred to as KP-HELLO),^38^ were kindly provided by Dr. Nikhil Joshi (Yale University). KP and KP-HELLO cells were cultured in RPMI1640 medium supplemented with 10% heat-inactivated FBS and 1% penicillin-streptomycin. Lewis lung carcinoma (LLC) cells (ATCC CRL-1642) and Vero E6 cells (ATCC CRL-1587) were maintained in DMEM with 10% heat-inactivated FBS and 1% penicillin-streptomycin. All the cells were cultured no more than 10 passages.

The SARS-CoV-2 mouse-adapted strain MA10 was passaged in Vero E6 cells and the titer was determined by plaque assay using Vero E6 cells. Influenza A/PR8/34 virus stock was generated in our laboratory.

#### Lung cancer models

For the orthotopic KP lung cancer model, 6- to 7-month-old male mice were infected with 5000 pfu of MA10 virus or PBS. In influenza A virus (IAV) experiments, 12-to 15-week-old female and male mice were infected with PR8 strain (100 pfu for females,120 pfu for males) or PBS. At 21 days post-infection, all mice were intratracheally inoculated with KP tumor cells (1×10^6^ cells in 50 ul PBS). Unless otherwise specified, lungs were harvested for analysis 3 weeks after tumor implantation. For CD8^+^ T cell functional assays, IAV infected mice were i.t. inoculated with 1×10^6^ KP-KELLO tumor cells. Three weeks later, mice were sacrificed, and total lung cells were prepared and stimulated ex vivo with 2 ug/ml GP33 peptide to determine cytokine-producing CD8^+^ T cells.

For the genetically engineered mouse model of lung cancer, KP mice were infected with 80 pfu of IAV or PBS. Three weeks later, tumor initiation was induced via intratracheal instillation of 2.5×10^8^ pfu Ad5-SPC-Cre virus (Viral Vector Core, University of Iowa), as previously described.^32,61^ Mice were sacrificed 12 weeks after tumor initiation or monitored for survival.

For the urethane-induced lung cancer model,^33^ young male mice were infected with IAV or PBS for 4 weeks prior to receiving weekly intraperitoneal injection of urethane (ethyl carbamate, EC; 1 mg per body weight) for 10 consecutive weeks. Mice were euthanized at 34 weeks after the first urethane dose, and lungs were collected for tumor burden analysis.

For subcutaneous KP tumor implantation, male mice were infected with PR8 for PBS for three weeks, then injected with 2×10^5^ KP tumor cells into the right flank. Tumor size was measured by calipers every three days, and tumor weights were determined at the endpoint. Tumor volume was calculated as follow: (length x width^2^)/2.

### METHOD DETAILS

#### Lung histology and Immunofluorescence

Lung tissues were harvested and fixed in 10% neutral-buffered formalin for a minimum of three days, and then sent to the Research Histology Core at UVA for processing, paraffin embedding, and sectioning (5 μm thickness). Hematoxylin and eosin (H&E) staining was performed with a standard method in the core. H&E slides were digitally scanned at 20X magnification in Biorepository and Tissue Research Facility (BTRF) at UVA. Tumor burden was quantified by outlining tumor regions and calculating the percentage of tumor area relative to total lung area using QuPath software. All analyses were performed in a blinded manner.

Immunofluorescence staining on paraffin sections was performed as previously described.^27^ Briefly, sections were deparaffinized in xylene and rehydrated through graded ethanol and subjected to heat-induced antigen retrieval using 1x Agilent Dako Target Retrieval Solution (pH 9, S236784) for 20 minutes. After PBS washes, sections were incubated overnight at 4°C with rabbit anti-Siglec5 (1:50, Abcam, ab320088) or anti-CD8a (1:300, Cell Signaling, #98941S) diluted in Dako Antibody Diluent. Following three PBS washes, sections were incubated for 2 hours at room temperature with goat anti-rabbit IgG Alexa Fluor 555 (1:300, ThermoFisher, A-21428). After additional PBS washes, sections were permeabilized with 0.5% Triton X-100 and 0.05% Tween-20 for 1 hour at room temperature, then incubated overnight at 4°C with Alexa Fluor 647-conjugated anti-myeloperoxidase (1:50, Abcam, ab252131). Nuclei were counterstained with DAPI (1:5000) for 3 minutes, and slides were mounted with ProLong Diamond Antifade Mountant. After curing for 24 hours, images were acquired using a 20× objective on an Olympus BX63 fluorescence microscope to quantify MPO⁺ neutrophils, MPO^+^ SiglecF⁺ neutrophils or CD8^+^ T cells in tumor areas. Image analysis was performed using OlyVIA and/or QuPath software. High-resolution images were obtained with a Zeiss LSM 880 confocal microscope using a 25x oil immersion objective (LD LCI Plan-Apochromat 25×/0.81 mm Korr DIC M27).

#### Tissue processing, flow cytometry, and FACS sorting

Mice were intravenously injected with 2 μg of anti-CD45 antibody diluted in 100 μl sterile PBS. Five minutes later, animals were euthanized for tissue collection. Bronchoalveolar lavage fluid (BALF) was collected by flushing the airway with a single 600 µl inoculum of FACS buffer (PBS + 2% FBS + 2 mM EDTA) via a tracheal incision. BALF was used for cytokines/chemokines analysis. Following BAL collection, the right ventricle was perfused with 10 ml chilled 1x PBS. The right lung lobes were harvested, minced, and digested with 180 U/ml collagenase type II (Worthington Biochemical) at 37 °C for 30 minutes. Tissue was disrupted using a gentleMACS tissue dissociator (Miltenyi), followed by red blood cell lysis, filtration through a 70 μm cell strainer, and resuspension in FACS buffer to obtain single-cell suspensions for flow cytometry.

For flow cytometric analysis, cells were stained with Zombie NIR live/dead dye and Fc receptor block for 10 minutes at room temperature, followed by incubation with antibodies against surface markers for 30 minutes on ice. Surface antibodies included: CD45, CD45.1, CD45.2, CD11b, CD11c, Ly6G, Ly6C, MHC II, CD64, MerTK, SiglecF, PD-L1, CD3, CD4, CD8, CD44, IFN-γ, TNFα. For intracellular staining, cells were first stimulated with 2 μg/ml GP33 peptide at 37 °C for 2 hours, followed by brefeldin A (1:1000) treatment for 4 hours prior to staining. Flow cytometry was performed using an Attune NxT cytometer (Invitrogen), and data were analyzed with FlowJo software.

For sorting of SiglecF^hi^ and SiglecF^lo^ neutrophils, tumor-bearing lungs from PBS- or IAV-infected mice were processed as above. Single-cell suspensions were enriched using anti-Ly6G microbeads (Miltenyi, 130-120-337), then stained with CD11b, Ly6G, and SiglecF antibodies. Cell sorting was performed using a BD Influx cell sorter in the UVA Flow Cytometry Core.

#### Parabiosis experiment

Age- and weight-matched female C57BL/6 congenic CD45.2 and CD45.1 mice were surgically joined in parabiosis as previously described.^62^ CD45.2 mouse was infected with IAV for 4 weeks before surgically jointed with PBS-treated CD45.1 mouse to form PBS-IAV pair, and PBS-PBS pair was included as control. For analgesia, carprofen (10 mg/kg) and buprenorphine (0.5 mg/kg) were administered intraperitoneally 1 hour prior to anesthesia with ketamine (100 mg/kg) and xylazine (10 mg/kg). Mice were shaved along the lateral trunk (elbow to knee), skin was sterilized with povidone-iodine and 70% ethanol, and matching longitudinal skin incisions were made. Olecranon and knee joints were aligned and sutured using 3-0 non-absorbable sutures, followed by continuous closure of the skin with 5-0 absorbable sutures both ventrally and dorsally. Animals received 0.5 ml sterile saline subcutaneously and triple antibiotic ointment was applied to the wound area postoperatively. Mice were placed on a warming pad until fully recovered, and DietGel food was provided on the cage floor to assist feeding. Post-surgical care included carprofen and buprenorphine for 48 hours, prophylactic enrofloxacin in drinking water (0.25 mg/ml) for one week, and daily monitoring for signs of distress. Three weeks after surgery, both mice were i.t. inoculated with KP tumor cells, and neutrophils and tumor burden were evaluated three weeks post-inoculation.

#### mRNA-Spike vaccination

SARS-CoV-2 mRNA-Spike vaccine from Moderna was purchased form Pharmacy at the University of Virginia and stored at -80C. For immunization, mice received two intramuscular injections of 1 μg mRNA-Spike formulated in 50 μl PBS, administered 21 days apart. Twenty-one days after the second dose, both mRNA-S- and vehicle-immunized mice were infected intranasally with either PBS or 5000 pfu of MA10 virus.

Mice body weight loss and survival were monitored daily. Three weeks after infection, all mice were intratracheally inoculated with 1 million KP tumor cells, and tumor burden was assessed three weeks later.

#### Bone marrow chimeras

Wild-type C57BL/6 mice were infected with PBS or IAV (n = 5 per group). At 21 days post-infection, bone marrow cells were isolated and transplanted into lethally irradiated (1100 rad) 12-week-old female naïve recipient mice to generate chimeric mice.

Following a 7-week reconstitution period, recipients were intratracheally inoculated with 1 million KP tumor cells, and tumor burden was assessed three weeks later.

#### In vivo tumor cell and neutrophil co-injection assay

A total of 2×10^5^ KP tumor cells were resuspended with an equal number of sorted SiglecF^hi^ or SiglecF^lo^ neutrophils in 100 ul of ice-cold 1x PBS and co-injected subcutaneously into the right flank of naïve recipient mice. To maintain neutrophil exposure within the tumor microenvironment, booster doses of matched SiglecF^hi^ or SiglecF^lo^ neutrophils were injected directly into the primary injection site on days 3 and 7 after initial implantation. Tumor growth was recorded weekly, and tumor were weighed at the endpoint (day 28).

#### Administration of antibodies and small molecule inhibitor

For prophylactic treatment, SCV2- or IAV-infected mice were randomly assigned to treatment groups one day prior to KP tumor cell intratracheal inoculation. Mice received intraperitoneal injections of anti-Ly6G antibody (200 μg/dose, every other day) or isotype control IgG, the CXCR2 inhibitor Reparixin (REP; 100 μg/dose, every other day) or vehicle control (DMSO), or intratracheal administration of anti-G-CSFR antibody (50 μg/dose, every 3 days) or control IgG. All treatments continued for the duration of the experiment.

For therapeutic treatment, SCV2-infected mice were divided into four groups one week after tumor implantation and treated with either DMSO plus IgG control, REP plus IgG control, anti-PD-L1 antibody (200 μg/dose, twice per week) plus DMSO, or a combination of REP and anti-PD-L1.

#### Multiplex cytokine/chemokine profiling

Cytokine and chemokine levels in BALF were quantified using a 32-plex cytokine array (Eve Technologies Inc.) according to the manufacturer’s instructions. Analytes with statistically significant differences between groups, determined by Mann–Whitney test, were selected and visualized in a heatmap generated using R. The level of TGF-b in BAL was measured by using TGF-beta 1 ELISA Kit (Biotechne, #DB100C) according to the manufacturer’s protocol.

#### In vitro restimulation assay

Lung single cell suspensions and AMs were prepared from SCV2-infected, IAV-infected, and their PBS-treated control mice. For total lung cell restimulation, 2×10⁵ cells were seeded in 24-well plates in RPMI-1640 supplemented with 10% FBS and 1% Penicillin/Streptomycin. Cells were stimulated overnight with vehicle, 20 ng/ml LPS, or equal numbers of UV-irradiated KP cells. UV-irradiated KP cells were prepared by exposing cells in PBS to UV-C light (254 nm) for 30 minutes in a tissue culture hood.

For AM restimulation, 1x10⁵ cells were seeded in 48-well plates and stimulated overnight with vehicle or 20 ng/ml LPS. Culture supernatants were collected for downstream ELISA analysis of G-CSF production by using Mouse G-CSF DuoSet ELISA kit (Biotechne, #DY414-05) following the manufacturer’s protocol.

#### Induction of SiglecF^+^ neutrophils

For in vitro induction of SiglecF^+^ neutrophils, bone marrow neutrophils were isolated form naïve mice using anti-Ly6G microbeads, achieving a purity exceeding 95% as confirmed by flow cytometry. Cells (1×10^5^ cells per well) were seeded in a 96-well plate and stimulated overnight with 10 ng/ml of recombinant G-CSF, GM-CSF, and TGF-β, or a combination of TGF-b with either G-CSF or GM-CSF. Following stimulation, cells were then washed and stained for SiglecF expression prior to flow cytometry analysis.

For in vivo SiglecF^+^ neutrophil induction, naïve mice were i.n. treated with 1μg rG-CSF and 1μg rTGF-β daily for three consecutive days. Vehicle was applied to control mice. On day 3, lungs were collected for flow cytometry analysis of SiglecF^+^ neutrophils.

#### Single cell RNA-Sequencing analysis

KP mice infected with IAV for 3 weeks were i.t. administered Ad5 SPC-Cre virus to initiate lung tumorigenesis. Lung single-cell suspensions were generated at weeks 6 (early-stage) and 12 (late-stage) after tumor induction for scRNA-Seq analysis. Tumor free PBS- and IAV-treated mice served as controls. Each group was representing pooled samples from 3-5 mice.

Single-cell libraries were prepared using the Chromium Single Cell 3′ Reagent Kit (10x Genomics) following the manufacturer’s protocol. Paired-end sequencing was performed on the DNBSEQ-G400 platform in rapid-run mode. Sequencing reads were aligned and quantified using the 10x Genomics Cell Ranger pipeline. Doublets were identified and removed using the *scDblFinder* R package. Data analysis was conducted using *Seurat* (v4.1.1), with quality filtering criteria of >200 detected genes, >1000 UMIs, and <5% mitochondrial gene content per cell. Downstream analysis included normalization, dimensionality reduction, clustering, and identification of cluster-specific marker genes and differentially expressed genes. Gene set enrichment analysis (GSEA) was performed using the *clusterProfiler* package based on marker genes identified by the *FindAllMarkers* function.

#### Single cell ATAC-Sequencing analysis

Lung single-cell suspensions freshly isolated from PBS- or SCV2-infected mice at 28 d.p.i. were processed for ATAC library construction using the Chromium Next GEM Single Cell ATAC Kit v2 (10x Genomics), following the manufacturer’s protocol. Libraries were sequenced on an Illumina NovaSeq S2 platform (50 bp paired-end, 100-cycle run). FASTQ files were generated and processed using 10x Genomics Cell Ranger ATAC (v2.1.0), with reads aligned to the mm10 reference genome. Approximately 8,000 nuclei were recovered per sample. Cells with fewer than 1,000 unique fragments or transcription start site (TSS) enrichment scores <5 were excluded. Fragment files were converted to a binarized cell-by-window matrix, retaining the top 20,000 most accessible peaks. Dimensionality reduction was performed using Latent Semantic Indexing (LSI), including TF-IDF transformation and singular value decomposition (SVD), followed by clustering with the Seurat R package (v3.1.4). Then peaks were recalled on each annotated cell type using *CallPeaks* function in *Signac* along with *MACS2*. Recalled peaks were combined together and further merged with the original peaks to form the final peak set. Peaks were annotated by *ChIPseeker* (v1.40.0) and *TxDb.Mmusculus.UCSC.mm10.knownGene* (v3.10.0). DNA sequence motif information was added to the chromatin assay based on *JASPAR2020* (v0.99.10) and *BSgenome.Mmusculus.UCSC.mm10* (v1.4.3), and motif activity was calculated using the *chromVAR* (v.1.26.0) implemented in *Signac*. Logistic regression implemented in *FindMarkers* function of *Seurat* was used to identify differential accessible peaks (DAPs) with library size of each cell treated as covariate. Note that only peaks detected in at least 5% cells in either group were included for differential analysis, and DAPs were defined as those with Benjamini & Hochberg (BH)-adjusted *P* value < 0.05. The Wilcoxon test was used to compare motif activity across groups, and motifs with BH-adjusted *P* < 0.05 and average difference of scaled activity score > 1 were defined as ones showing differential activity. In addition, a hypergeometric test was used to identify motifs overrepresented in DAPs for target cell types compared to 50,000 background peaks matched for GC content.

SCENIC+ workflow (v.1.0a1) was used to infer gene regulation networks (GRNs) by integrating scRNA-seq and scATAC-seq data. In brief, *pycisTopic* (v.2.0a0) was first used to define candidate transcription factor-binding regions from scATAC-seq data.

Then, *pycisTarget* (v 1.0a2), together with its built-in motif collection, was used to perform motif enrichment analysis and construct custom cisTarget database. Cluster-Buster was also used to score and rank the sequence of consensus peaks for each motif in collection and generate cisTarget regions vs motifs database. Finally, scRNA-seq and scATAC-seq data were integrated using *SCENIC*+ to infer TF-(motif)-region accessibility-gene expression relationship, where GRNBoost2 (*Arboreto*, v.0.1.6) was used to quantify the importance and direction of TFs to target genes and regions to target genes, and eRegulon activity scores were computed with AUCell based on target gene expression and target region accessibility. The Wilcoxon test was used to identify motifs with differential activity across groups. Those with BH-adjusted *P* < 0.05 and average difference of activity score > 0.01.

#### Survival Analysis of Lung Adenocarcinoma in TCGA

Survival analysis was performed using publicly available TCGA LUAD (lung adenocarcinoma) transcriptomic and clinical data accessed through the TCGAbiolinks and survival R packages. Expression levels of the top 100 differentially expressed genes were used to generate gene signature scores, based on comparisons between SiglecF^hi^ neutrophils from IAV-infected versus PBS-treated mice. Additionally, a set of conserved genes shared between mouse SiglecF^hi^ neutrophils and human SiglecF^hi^-like neutrophils.^24^ For each patient, a signature score was calculated as the average normalized expression of the selected genes. Patients were stratified into “high” and “low” expression groups using the median or percentile-based threshold that maximized survival discrimination. Kaplan–Meier plots and log-rank tests were generated using the survfit function, and univariate Cox proportional hazards regression was used to evaluate statistical significance.

### QUANTIFICATION AND STATISTICAL ANALYSIS

Statistical analysis was performed using GraphPad Prism (version 9, GraphPad Software). For all experiments with two groups, two-tailed Mann-Whitney tests were used. One-way ANOVA with correction for multiple comparisons was performed for experiments with more than two groups. Two-way ANOVA was used for tumor kinetic experiment. Log-rank Mantel-Cox test was used for survival analysis. The data was presented as mean ± SEM. Figure legends specify the statistical analysis used.

## SUPPLEMENTARY FIGURE LEGENDS

**Figure S1.**
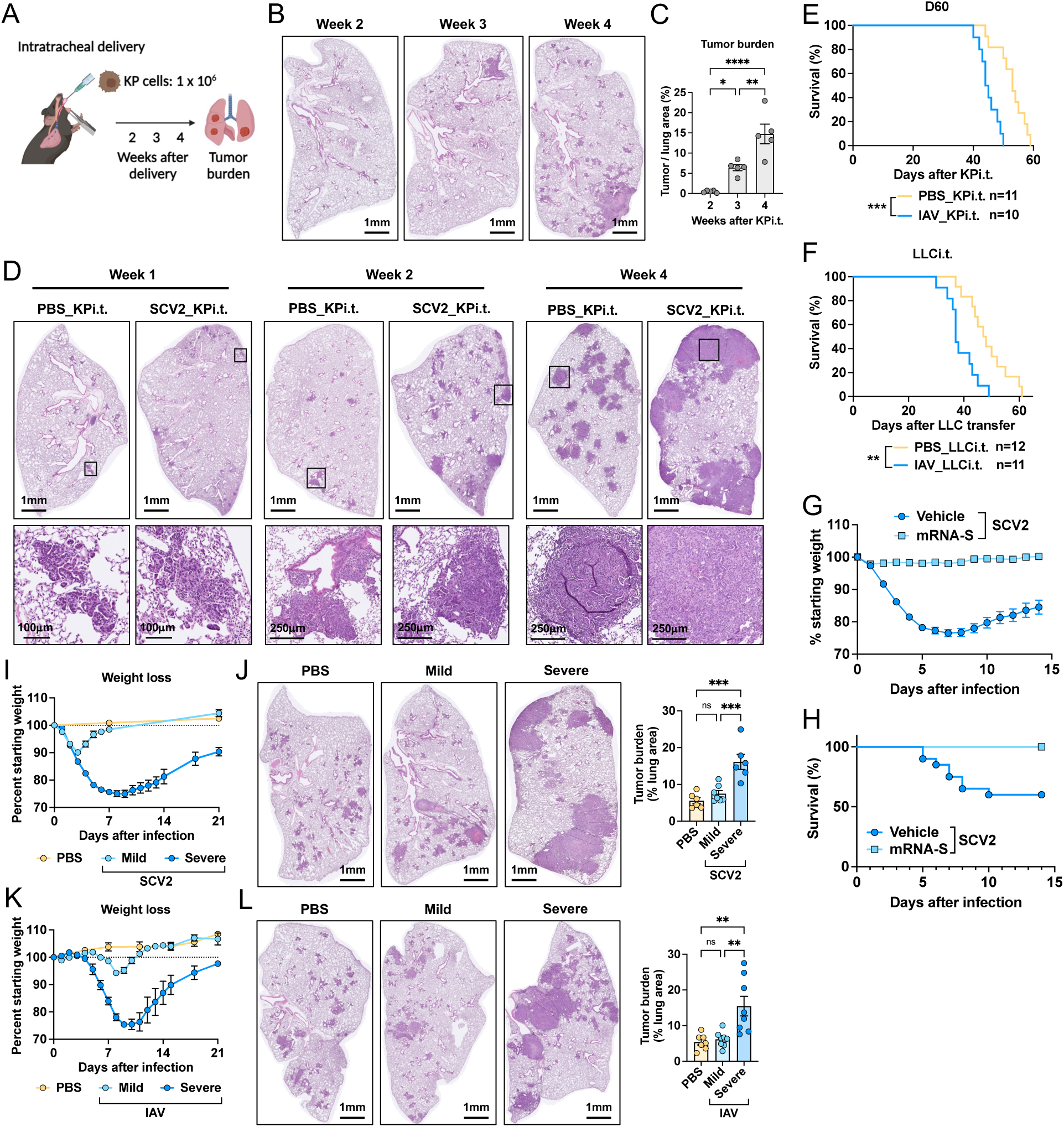
Prior respiratory viral infections prime enhanced lung tumor growth, related to Figure 1. (**A**) schematic showing intratracheal (i.t.) delivery of KP tumor cells and timeline for tumor burden measurement. (**B and C**) Representative image (B) and quantification of tumor burden (C) in naive mice from weeks 2 to 4 after tumor cell inoculation. (D) Representative images of tumor-bearing lungs from SCV2-Infected and PBS-treated mice at weeks 1, 2, and 4 after inoculation. Outlines indicate roomed-in tumors. (E) Survival curves of tumor-bearing mice from PBS-treated and IAV-infected (D60) groups. (F) Survival curves of mice from PBS-treated and IAV-infected (D21) groups following i.t. inoculation of LLC tumor cells. (**G and H**) Body weight change (G) and mortality rate (H) in mice immunized with two doses of mRNA-S vaccine or PBS following SCV2 infection. (I) Weight loss in naïve mice following mild (50 pfu) and severe (5000 pfu) SCV2 infection. (J) Representative lung tumors (left) and tumor burden quantification (right) at 3 weeks post-tumor initiation in mildly and severely SCV2-infected or PBS-treated mice. (K) Weight loss in naïve mice following mild (5 pfu) and severe (100 pfu) IAV. (L) Representative lung tumors (left) and tumor burden quantification (right) at 3 weeks post-tumor inoculation in mildly and severely IAV-infected or PBS-treated mice. Data represent two independent experiments or are pooled from two experiments (E-L). Graphs display mean ± SEM. Statistical significance was assessed by one-way ANOVA (C, J and L) or Log-rank Mantel-Cox test (E and F). (ns *p* > 0.05, \**p* < 0.05, \*\**p* < 0.01, \*\*\**p* < 0.001, \*\*\*\**p* < 0.0001)

**Figure S2.**
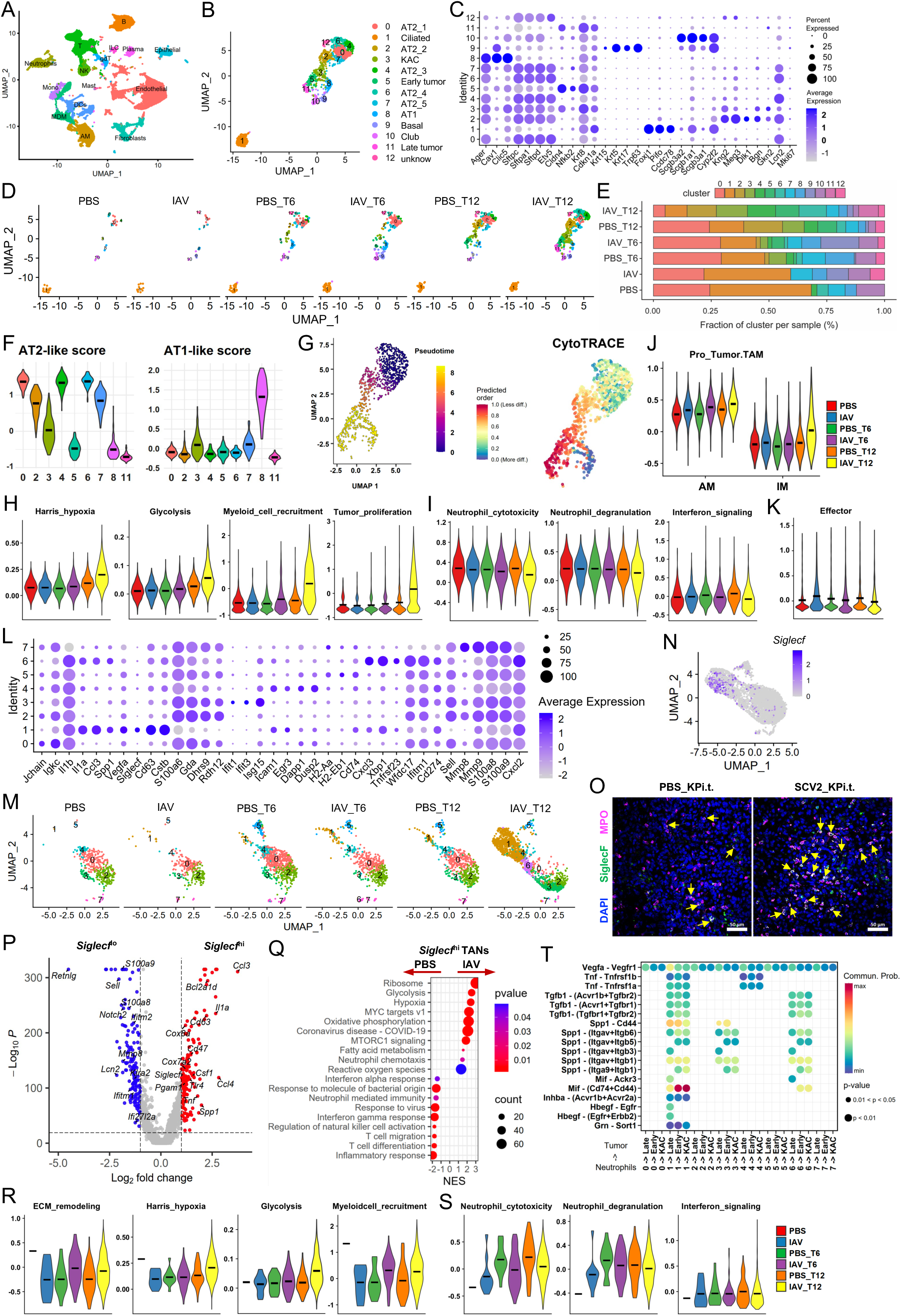
Single-cell characterization of tumor-bearing lungs, related to Figure 2. (A) UMAP plots of whole lung cells from 6 study groups. (B) UMAP distribution of epithelial cell subsets. (C) Dot heatmap showing the expression of typical marker genes for epithelial cell types. (D) UMAP distribution of epithelial cell subsets stratified by groups. (E) Stacked bar plot showing the relative proportions of 13 epithelial cell subsets across the six groups. (F) Violin plots displaying AT2-like and AT1-like gene signature scores across 9 alveolar epithelial clusters. (G) Trajectories of alveolar and tumor cells colored by inferred pseudotime and CytoTRACE scores. (H) Neutrophil pro-tumor signature scores across experimental groups. (I) Neutrophil anti-tumor signature scores across experimental groups (J) Pro-tumor TAM signature scores in AMs and IMs among 6 groups. (K) CD8^+^ T cell effector function scores across 6 groups. (L) Dot heatmap showing the expression of typical marker genes in neutrophil subsets. (M) UMAP distribution of neutrophil subsets across groups. (N) Feature plot showing *Siglecf* expression in neutrophil clusters. (O) Representative immunofluorescence images showing co-localization of MPO and SiglecF in tumors from PBS_KPi.t. and SCV2_KPi.t. groups. Scale bar, 50 μm. (P) A volcano plot visualizes DEGs between *Siglecf*^hi^ and *Siglecf*^lo^ neutrophils, highlighting genes with FDR < 0.05 and Log2 FC > 1 in blue (down-regulated) and red (up-regulated). (Q) Pathways enrichment analysis (Hallmark and GO database) of DEGs between IAV-*Siglecf*^hi^ and PBS-*Siglecf*^lo^ neutrophils. Cutoffs: p.adj < 0.05; NES > 1. (R) Pro-tumor neutrophil signature scores in *Siglecf*^hi^ TANs across experimental groups. (S) Anti-tumor neutrophil signature scores in *Siglecf*^hi^ TANs across experimental groups. (T) Dot plot showing CellChat-inferred intercellular signaling from neutrophil clusters (sender) to tumor clusters (receivers). Dot size represents p-value and color indicates communication probability.

**Figure S3.**
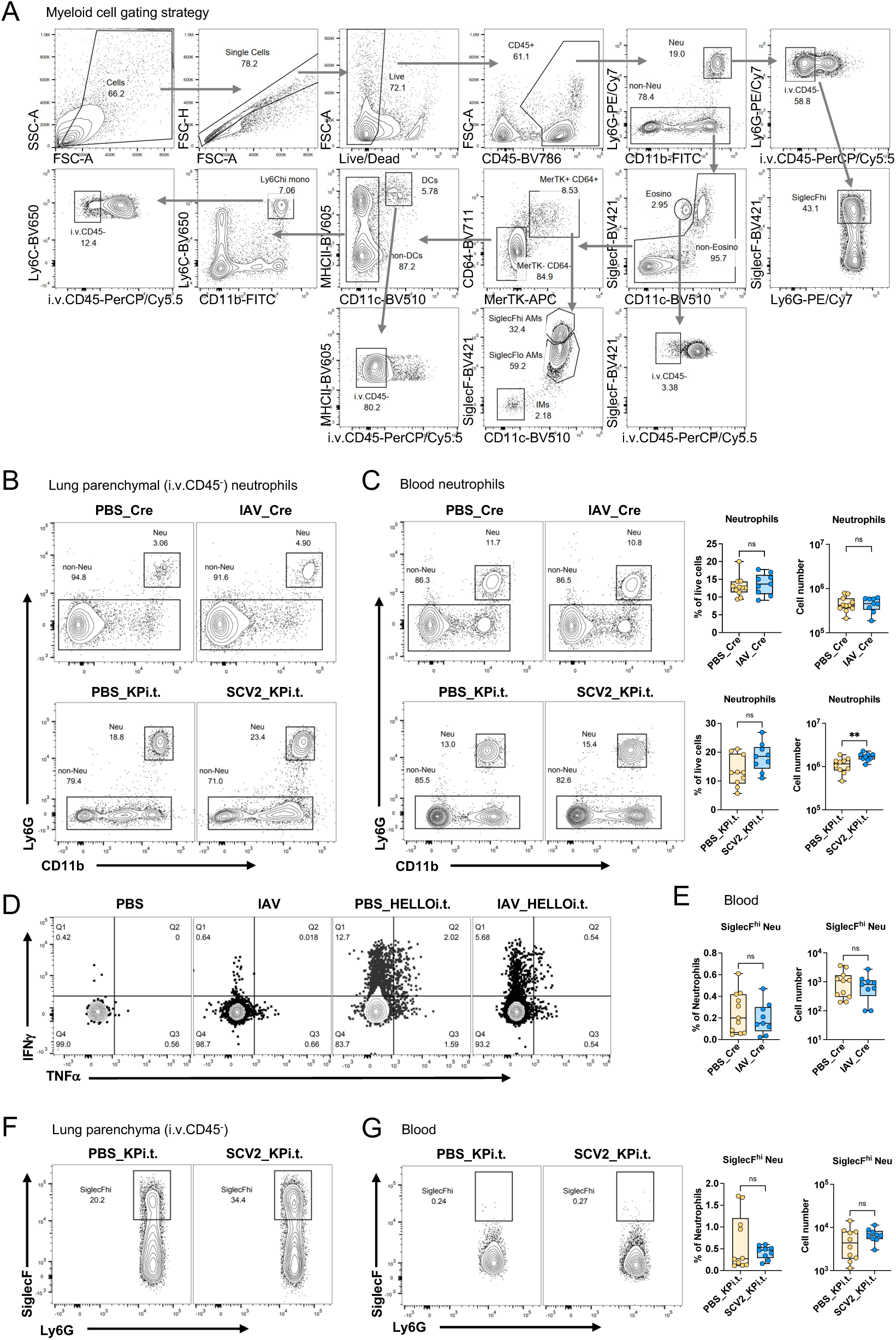
FACS gating strategy and neutrophil analysis, related to Figure 2. (A) Representative flow cytometry plots illustrating gating strategy for myeloid cells in lungs. (B) Flow cytometry plots showing lung parenchymal (i.v.CD45^-^) neutrophils in IAV_Cre and SCV2_KPi.t versus their respective PBS-treated controls. (C) Flow cytometry plots and quantification of blood neutrophils in IAV_Cre and SCV2_KPi.t. mice and their respective PBS-treated mice, including relative percentages and total cell counts. (D) Representative flow cytometry plots showing lung parenchymal (i.v.CD45^-^) CD8^+^ T cells producing IFNγ and TNFα following6-hour GP33 peptide stimulation in IAV_KP-HELLOi.t. and PBS-treated tumor mice, three weeks after tumor cell inoculation. (E) The proportion and total cell count of SiglecF^hi^ neutrophils in the blood of IAV_Cre and PBS_Cre mice. (F) Flow cytometry plots of lung parenchymal (i.v.CD45^-^) SiglecF^hi^ neutrophils in SCV2_KPi.t. and PBS-treated mice. (G) Flow cytometry plots and quantification of SiglecF^hi^ neutrophils in the blood of SCV2_KPi.t. and PBS-treated mice, including relative frequencies and cell counts. Data are represented of two independent experiments or pooled from two (C bottom, and G) or three experiments (C top, and E). Graphs show mean ± SEM. Statistical significance was assessed by Mann-Whitney test. (ns *p* > 0.05, \*\**p* < 0.01)

**Figure S4.**
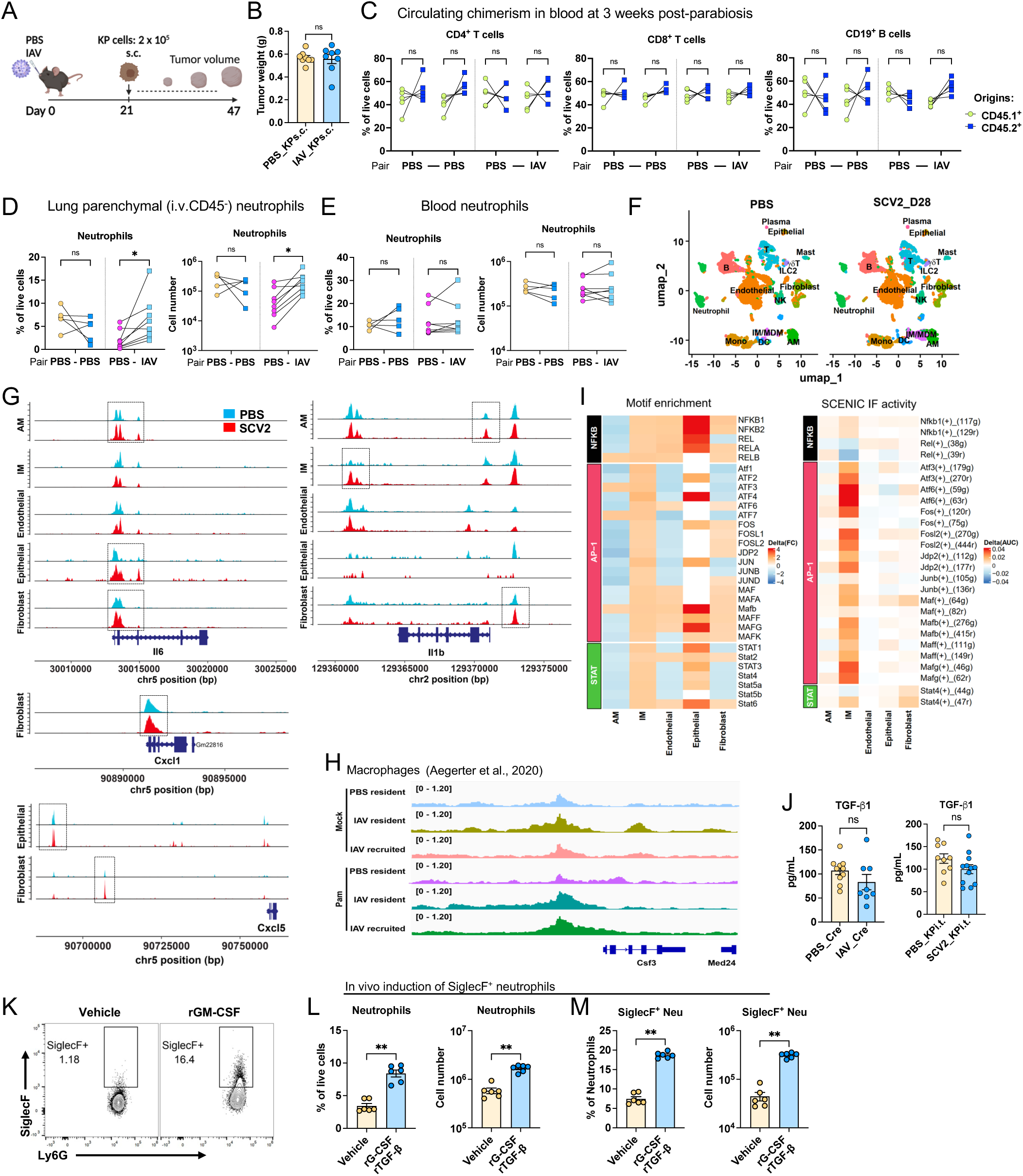
Viral infection epigenetically imprints the lung to promote tumor growth, related to Figure 3. (A) Model of KP tumor cells s.c. implantation at 21 days after IAV infection. (B) Tumor weight from (A) measured 26 days after implantation. (C) Examination of circulating chimerism in blood 3 weeks after parabiosis. Frequencies of CD4^+^ T, CD8^+^ T and CD19^+^ B cells were shown. (D) Proportion and total cell counts of neutrophils in the lung parenchyma from parabiosis experiments. (E) Proportion and total cell counts of neutrophils in the blood from parabiosis experiments. (F) UMAP plots of whole lung cells from scATAC-Seq of SCV2-infected and PBS-treated mice at 28 d.p.i.. (G) scATAC-seq data showing accessible regions within the *Csf3*, *Il6*, *Il1b*, *Cxcl1* and *Cxcl5* loci of the indicated lung cell types. Dashed box highlighted peaks increased in SCV2 groups compared to PBS groups. (H) IGV tracks of ATAC-seq data from AMs showing the *Csf3* locus and upstream peaks. (I) Motif enrichment and SCENIC-inferred IF activity analysis, showing activation of NF-kB, AP-1 and STAT family TFs. (J) TGF-β levels in BAL from IAV_Cre mice (week 12) and SCV2_KPi.t. mice (week 3), compared to their respective PBS-treated controls. (K) Flow cytometry dot plots showing the induction of SiglecF^+^ neutrophils after rGM-CSF treatment. (**L and M**) In vivo induction of SiglecF^+^ neutrophils in lungs of mice treated intranasally with rG-CSF and rTGF-β for three days. Flow cytometry plots and percentage of total neutrophils (K) and SiglecF^+^ neutrophils (L) in lung tissue. Data represent two independent experiments or pooled from two experiments (B, D, E, and J). Graphs display mean ± SEM. Statistical significance was assessed by Mann-Whitney test (B, J, L and M) and two-way ANOVA (C, D and E). (ns *p* > 0.05, \**p* < 0.05, \*\**p* < 0.01)

**Figure S5.**
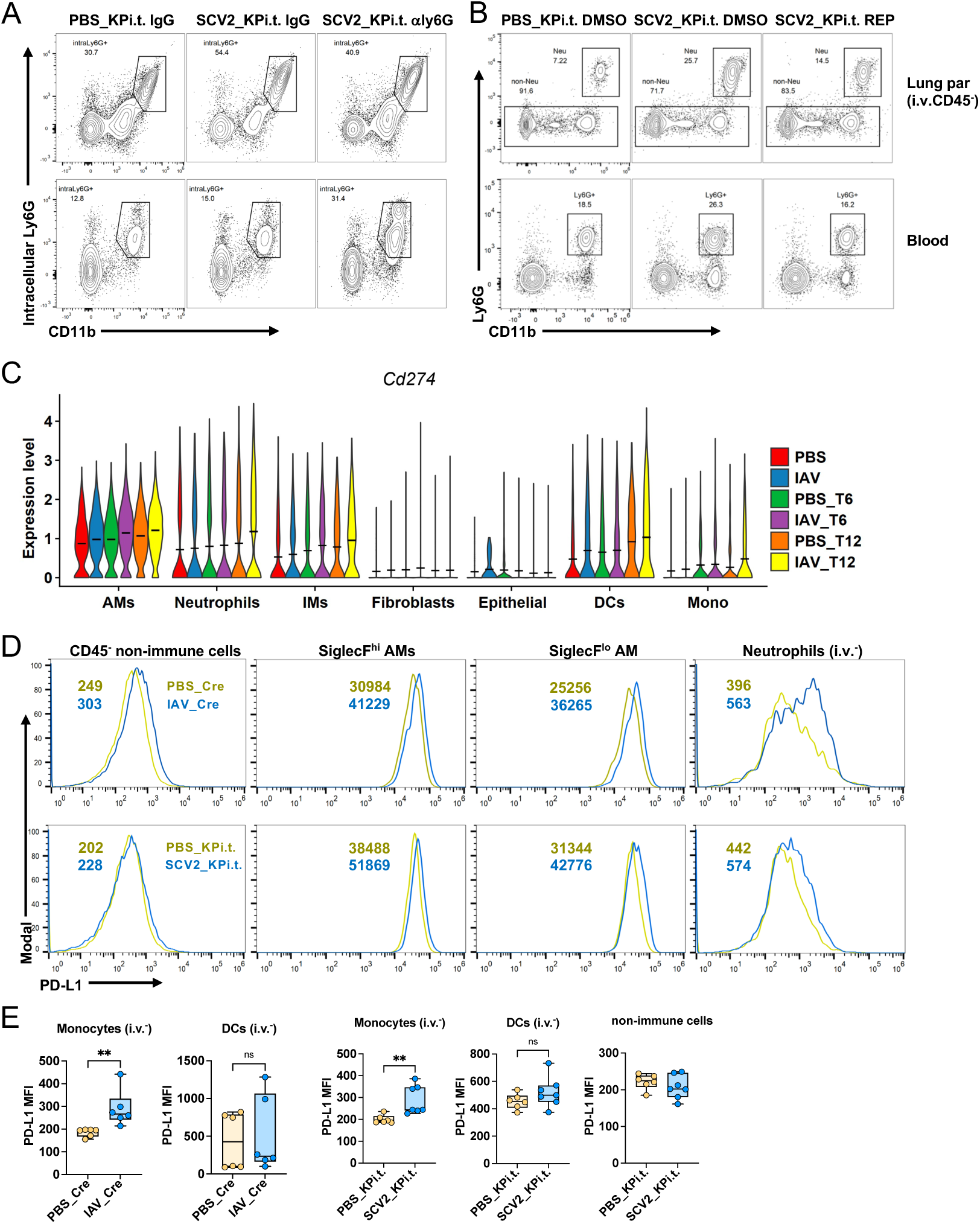
Neutrophil depletion efficiency and PD-L1 expression, related to Figure 4. (**A and B**) Flow cytometry plots showing neutrophil depletion in the SCV2_KPi.t. model following treatment with anti-Ly6G antibody (A) or CXCR2 inhibitor Reparixin (B). (**C**) *Cd274* expression across various cell types in 6 experimental groups, as determined by scRNA-Seq. (**D and E**) PD-L1 expression in multiple lung cells from IAV_Cre and SCV2_KPi.t. mice, shown as representative histograms (D) and quantified MFI (E). Data represent two independent experiments or are pooled from two experiments (E). Graphs display mean ± SEM. Statistical significance was assessed by Mann-Whitney test (E) and two-way ANOVA (C, D and E). (ns *p* > 0.05, \*\**p* < 0.01)

---

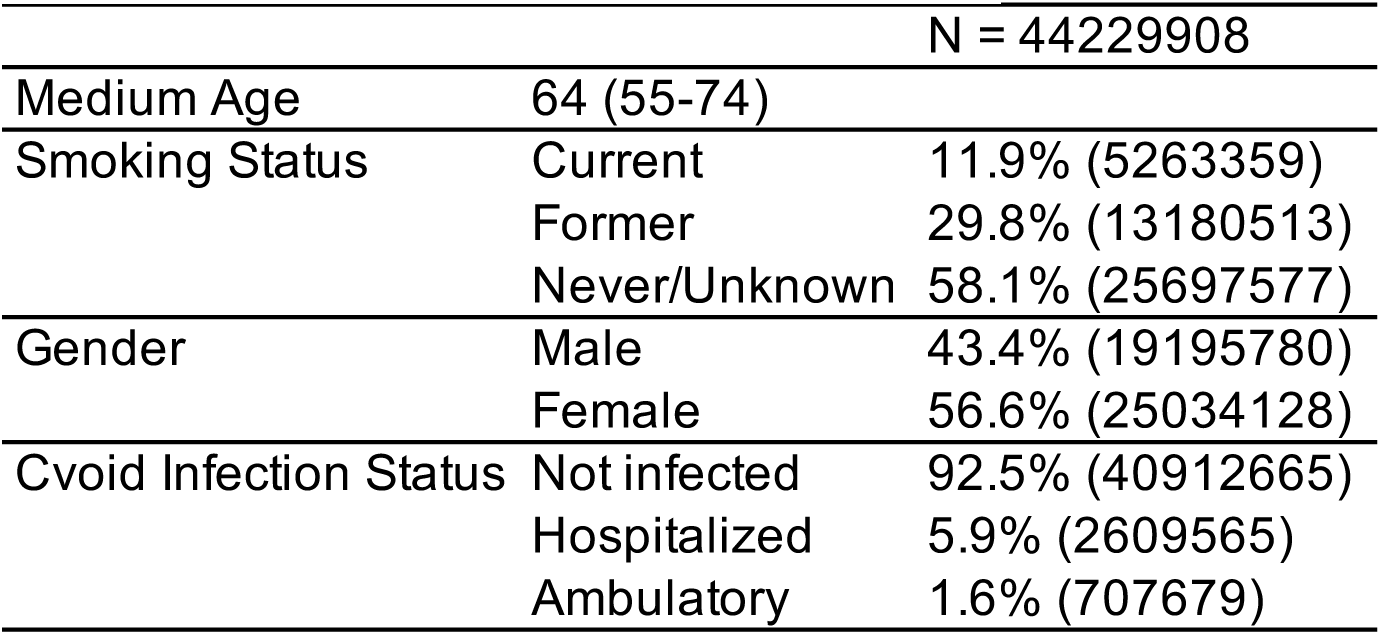
Cohort demographic information.

|  |  |
| --- | --- |
| AT1-like | AT2-like |
| Ager | Fos |
| Emp2 | Egr1 |
| Cav1 | Zfp36L1 |
| Hopx | Klf6 |
| Clic5 | Sftpb |
|  | Lrrk2 |
|  | Scgb3A2 |
|  | Sftpa2 |
|  | Sftpc |
|  | Sftpa1 |

| Immunosupp | ECMremodel | Tumorprolife | Myeloidcellr | Angiogenesis | Neutrophilde | Neutrophilcy |
| --- | --- | --- | --- | --- | --- | --- |
| Havcr2 | Adamdec1 | Tgfb1 | Ccl3 | Vegfa | Abr | Ncf1 |
| Fcgr2b | Ctsc | Tnf | Mif | Snd1 | Anxa3 | Myd88 |
| Il4ra | Ctsb | Il1a | Cxcl14 | Mtdh | Cd177 | Trem3 |
| Cd274 | Rgcc |  | Csf1 | Itga5 | Itgam | Trem1 |
| Hif1a | Ctss |  | Vegfa | Tnf | Itgb2 | Tusc2 |
|  | Ctsz |  | Ccl4 | Cxcl3 | Itgb2l | Cybb |
|  | Adam17 |  | Cxcl3 | Anxa3 | Pikfyve | Cybc1 |
|  | Adam10 |  |  | Hmgb1 | Pram1 | Ncf2 |
|  | Adam8 |  |  | Hif1a | Ptafr | Ncf4 |
|  |  |  |  | Sema4d | Spi1 | Rac1 |
|  |  |  |  | Lrg1 | Stx11 | Rac2 |
|  |  |  |  | Chil1 | Syk |  |

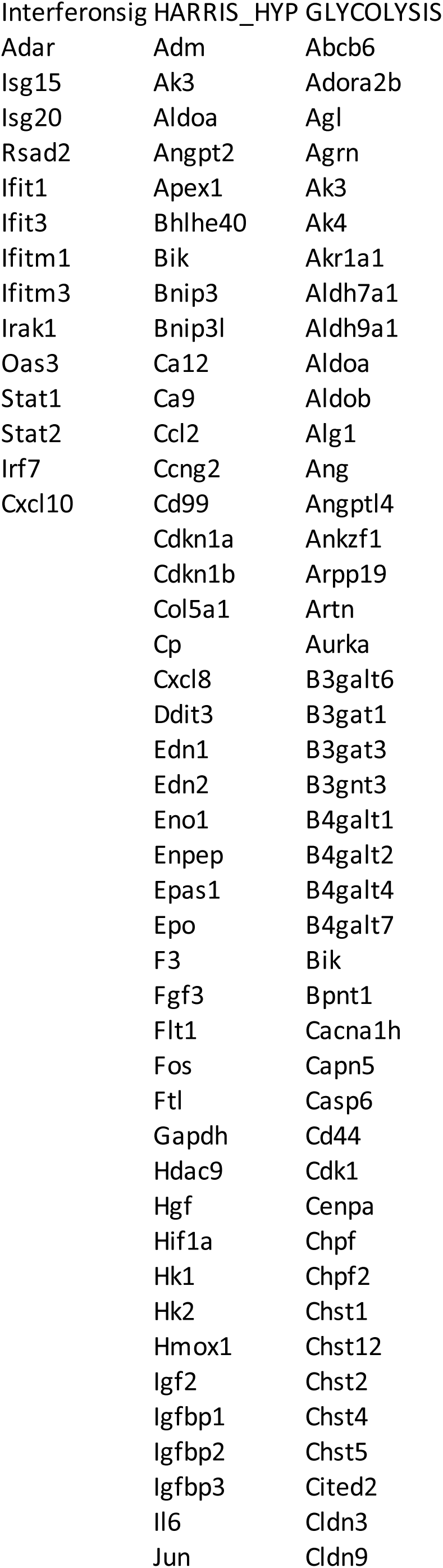

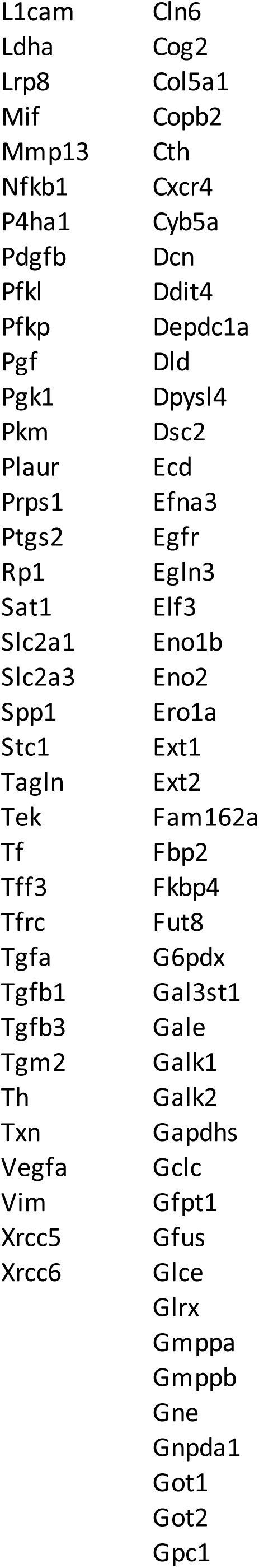

|  |  |
| --- | --- |
| L1cam | Cln6 |
| Ldha | Cog2 |
| Lrp8 | Col5a1 |
| Mif | Copb2 |
| Mmp13 | Cth |
| Nfkb1 | Cxcr4 |
| P4ha1 | Cyb5a |
| Pdgfb | Dcn |
| Pfkl | Ddit4 |
| Pfkp | Depdc1a |
| Pgf | Dld |
| Pgk1 | Dpysl4 |
| Pkm | Dsc2 |
| Plaur | Ecd |
| Prps1 | Efna3 |
| Ptgs2 | Egfr |
| Rp1 | Egln3 |
| Sat1 | Elf3 |
| Slc2a1 | Eno1b |
| Slc2a3 | Eno2 |
| Spp1 | Ero1a |
| Stc1 | Ext1 |
| Tagln | Ext2 |
| Tek | Fam162a |
| Tf | Fbp2 |
| Tff3 | Fkbp4 |
| Tfrc | Fut8 |
| Tgfa | G6pdx |
| Tgfb1 | Gal3st1 |
| Tgfb3 | Gale |
| Tgm2 | Galk1 |
| Th | Galk2 |
| Txn | Gapdhs |
| Vegfa | Gclc |
| Vim | Gfpt1 |
| Xrcc5 | Gfus |
| Xrcc6 | Glce |
|  | Glrx |
|  | Gmppa |
|  | Gmppb |
|  | Gne |
|  | Gnpda1 |
|  | Got1 |
|  | Got2 |
|  | Gpc1 |

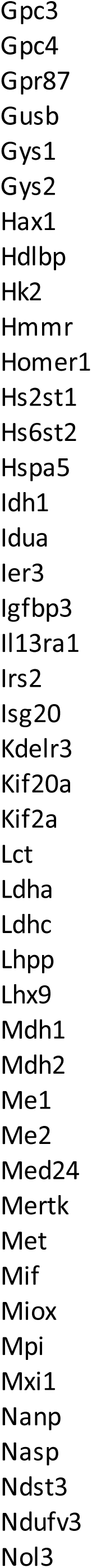

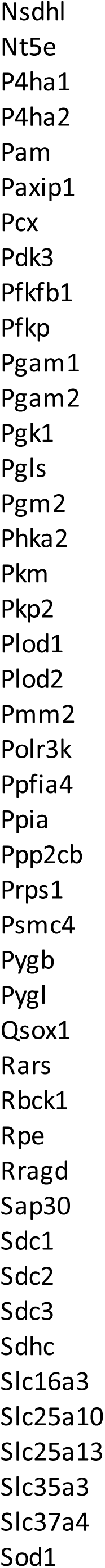

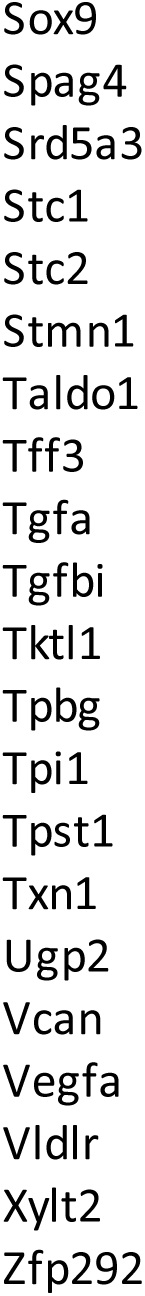

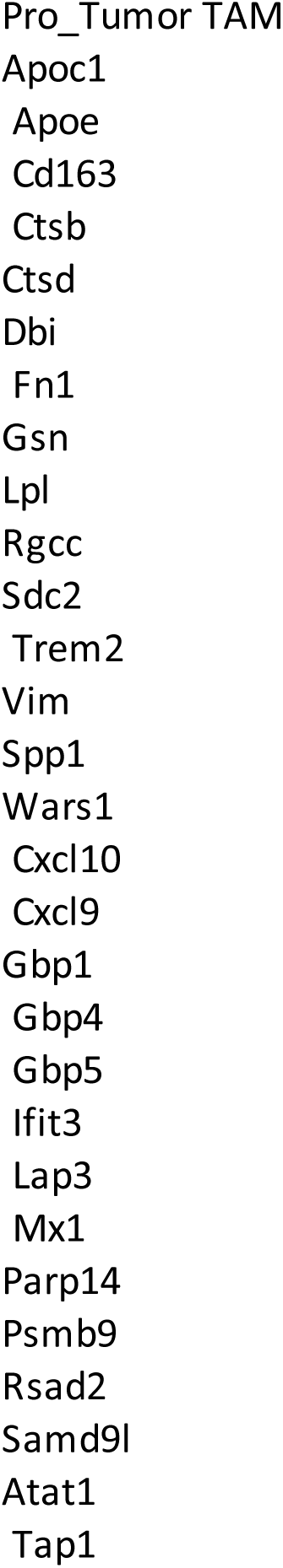

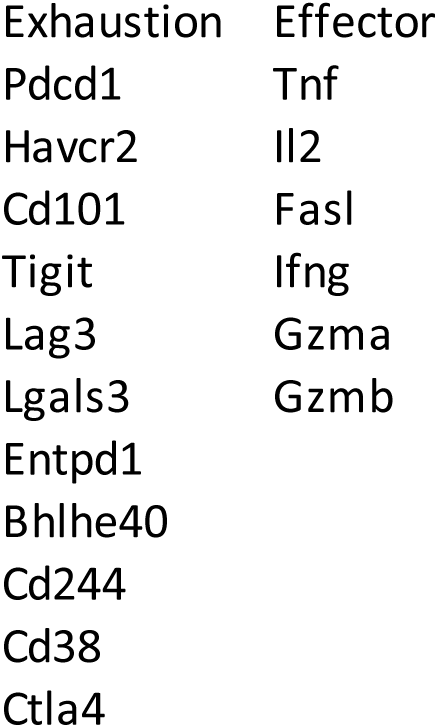

| Exhaustion | Effector |
| --- | --- |
| Pdcd1 | Tnf |
| Havcr2 | Il2 |
| Cd101 | Fasl |
| Tigit | Ifng |
| Lag3 | Gzma |
| Lgals3 | Gzmb |
| Entpd1 |  |
| Bhlhe40 |  |
| Cd244 |  |
| Cd38 |  |
| Ctla4 |  |

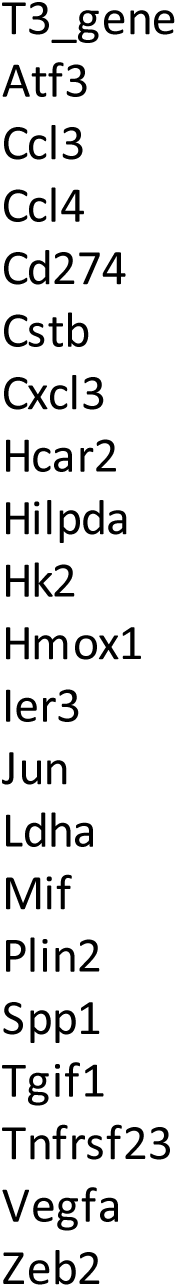

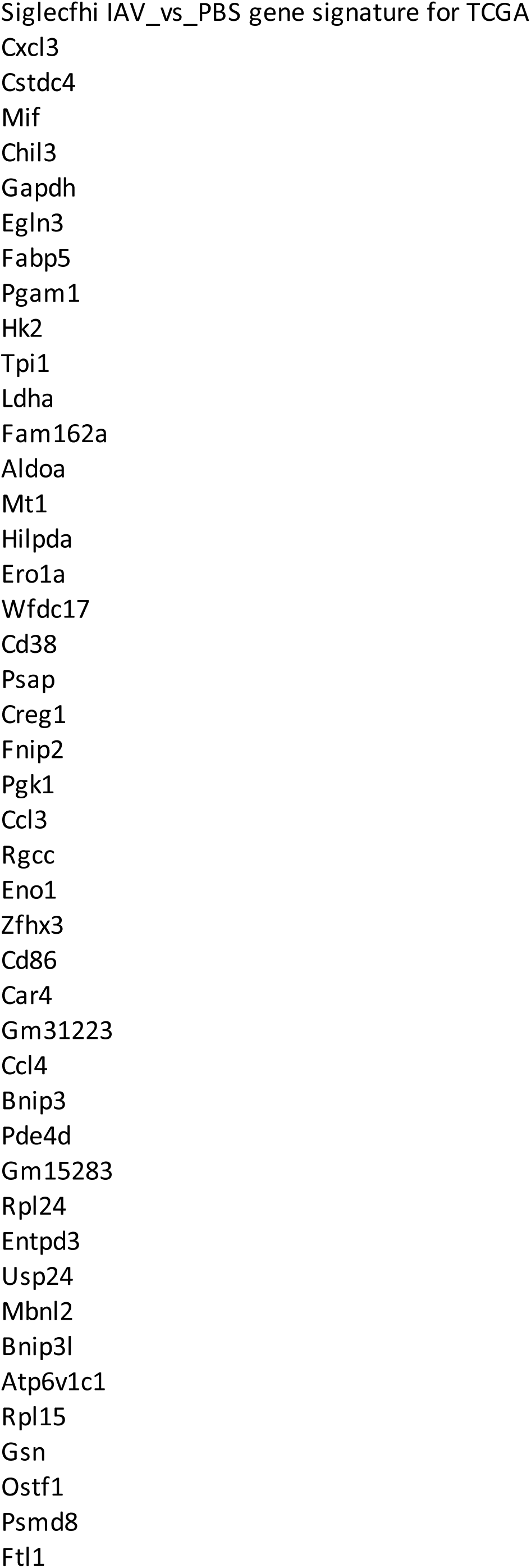

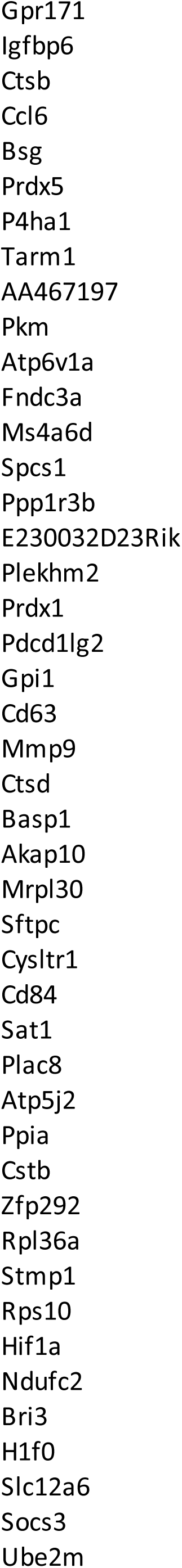

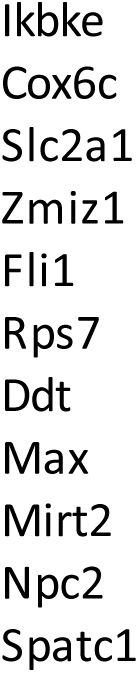

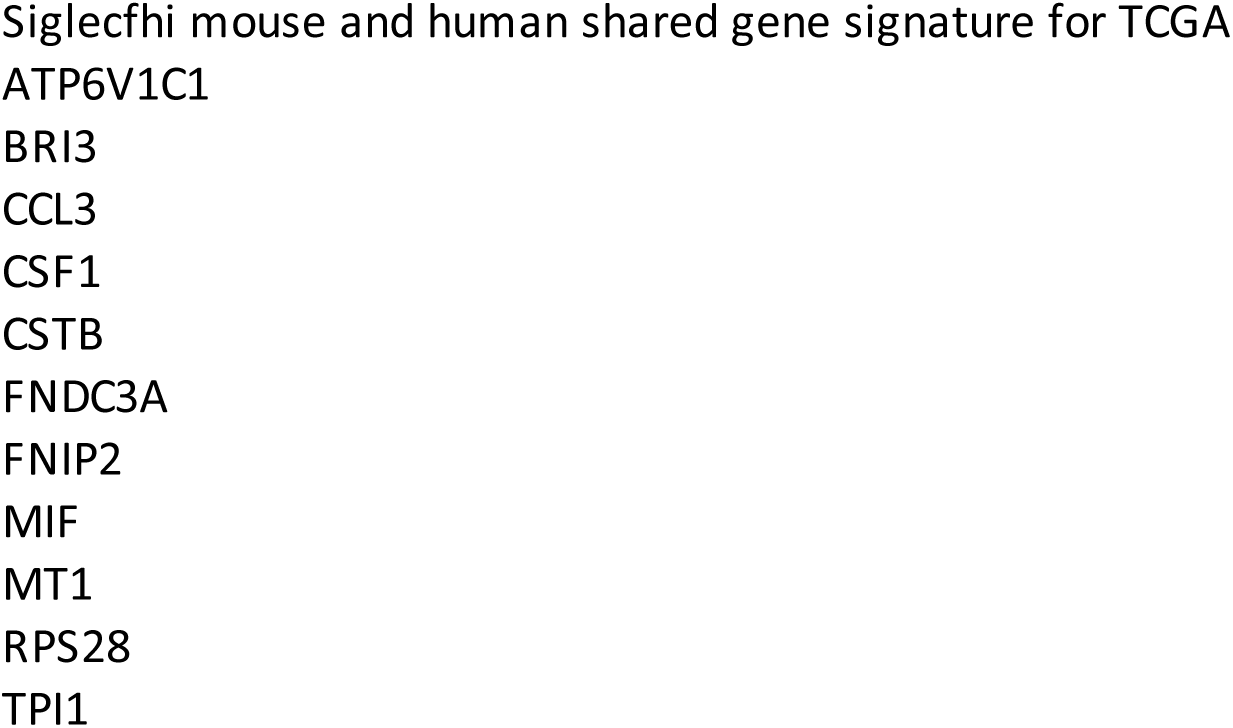

## REFERENCES

1. Al-Aly, Z., Davis, H., McCorkell, L., Soares, L., Wulf-Hanson, S., Iwasaki, A., and Topol, E.J. (2024). Long COVID science, research and policy. Nat Med 30, 2148–2164. 10.1038/s41591-024-03173-6.

2. Mehandru, S., and Merad, M. (2022). Pathological sequelae of long-haul COVID. Nat Immunol 23, 194–202. 10.1038/s41590-021-01104-y.

3. Nalbandian, A., Sehgal, K., Gupta, A., Madhavan, M.V., McGroder, C., Stevens, J.S., Cook, J.R., Nordvig, A.S., Shalev, D., Sehrawat, T.S., et al. (2021). Post-acute COVID-19 syndrome. Nat Med 27, 601–615. 10.1038/s41591-021-01283-z.

4. Choutka, J., Jansari, V., Hornig, M., and Iwasaki, A. (2022). Unexplained post-acute infection syndromes. Nat Med 28, 911–923. 10.1038/s41591-022-01810-6.

5. Wei, X., Narasimhan, H., Zhu, B., and Sun, J. (2023). Host Recovery from Respiratory Viral Infection. Annu Rev Immunol 41, 277–300. 10.1146/annurev-immunol-101921-040450.

6. Narasimhan, H., Wu, Y., Goplen, N.P., and Sun, J. (2022). Immune determinants of chronic sequelae after respiratory viral infection. Sci Immunol 7, eabm7996. 10.1126/sciimmunol.abm7996.

7. Xie, Y., Choi, T., and Al-Aly, Z. (2024). Long-term outcomes following hospital admission for COVID-19 versus seasonal influenza: a cohort study. Lancet Infect Dis 24, 239–255. 10.1016/S1473-3099(23)00684-9.

8. Weng, C.F., Chen, L.J., Lin, C.W., Chen, H.M., Lee, H.H., Ling, T.Y., and Hsiao, F.Y. (2019). Association between the risk of lung cancer and influenza: A population-based nested case-control study. Int J Infect Dis 88, 8–13. 10.1016/j.ijid.2019.07.030.

9. Meng, M., Wei, R., Wu, Y., Zeng, R., Luo, D., Ma, Y., Zhang, L., Huang, W., Zeng, H., Leung, F.W., et al. (2024). Long-term risks of respiratory diseases in patients infected with SARS-CoV-2: a longitudinal, population-based cohort study. EClinicalMedicine 69, 102500. 10.1016/j.eclinm.2024.102500.

10. Greten, F.R., and Grivennikov, S.I. (2019). Inflammation and Cancer: Triggers, Mechanisms, and Consequences. Immunity 51, 27–41. 10.1016/j.immuni.2019.06.025.

11. Denk, D., and Greten, F.R. (2022). Inflammation: the incubator of the tumor microenvironment. Trends Cancer 8, 901–914. 10.1016/j.trecan.2022.07.002.

12. Gagiannis, D., Hackenbroch, C., Bloch, W., Zech, F., Kirchhoff, F., Djudjaj, S., von Stillfried, S., Bulow, R., Boor, P., and Steinestel, K. (2023). Clinical, Imaging, and Histopathological Features of Pulmonary Sequelae after Mild COVID-19. Am J Respir Crit Care Med 208, 618–621. 10.1164/rccm.202302-0285LE.

13. Schultheiss, C., Willscher, E., Paschold, L., Gottschick, C., Klee, B., Henkes, S.S., Bosurgi, L., Dutzmann, J., Sedding, D., Frese, T., et al. (2022). The IL-1beta, IL-6, and TNF cytokine triad is associated with post-acute sequelae of COVID-19. Cell Rep Med 3, 100663. 10.1016/j.xcrm.2022.100663.

14. Canderan, G., Muehling, L.M., Kadl, A., Ladd, S., Bonham, C., Cross, C.E., Lima, S.M., Yin, X., Sturek, J.M., Wilson, J.M., et al. (2025). Distinct type 1 immune networks underlie the severity of restrictive lung disease after COVID-19. Nat Immunol 26, 595–606. 10.1038/s41590-025-02110-0.

15. Kay, J., Thadhani, E., Samson, L., and Engelward, B. (2019). Inflammation-induced DNA damage, mutations and cancer. DNA Repair (Amst) 83, 102673. 10.1016/j.dnarep.2019.102673.

16. Jaiswal, A., Shrivastav, S., Kushwaha, H.R., Chaturvedi, R., and Singh, R.P. (2024). Oncogenic potential of SARS-CoV-2-targeting hallmarks of cancer pathways. Cell Commun Signal 22, 447. 10.1186/s12964-024-01818-0.

17. Weeden, C.E., Hill, W., Lim, E.L., Gronroos, E., and Swanton, C. (2023). Impact of risk factors on early cancer evolution. Cell 186, 1541–1563. 10.1016/j.cell.2023.03.013.

18. Hedrick, C.C., and Malanchi, I. (2022). Neutrophils in cancer: heterogeneous and multifaceted. Nat Rev Immunol 22, 173–187. 10.1038/s41577-021-00571-6.

19. Ng, M., Cerezo-Wallis, D., Ng, L.G., and Hidalgo, A. (2025). Adaptations of neutrophils in cancer. Immunity 58, 40–58. 10.1016/j.immuni.2024.12.009.

20. Eruslanov, E., Nefedova, Y., and Gabrilovich, D.I. (2025). The heterogeneity of neutrophils in cancer and its implication for therapeutic targeting. Nat Immunol 26, 17–28. 10.1038/s41590-024-02029-y.

21. Ng, M.S.F., Kwok, I., Tan, L., Shi, C., Cerezo-Wallis, D., Tan, Y., Leong, K., Calvo, G.F., Yang, K., Zhang, Y., et al. (2024). Deterministic reprogramming of neutrophils within tumors. Science 383, eadf6493. 10.1126/science.adf6493.

22. Xue, R., Zhang, Q., Cao, Q., Kong, R., Xiang, X., Liu, H., Feng, M., Wang, F., Cheng, J., Li, Z., et al. (2022). Liver tumour immune microenvironment subtypes and neutrophil heterogeneity. Nature 612, 141–147. 10.1038/s41586-022-05400-x.

23. Salcher, S., Sturm, G., Horvath, L., Untergasser, G., Kuempers, C., Fotakis, G., Panizzolo, E., Martowicz, A., Trebo, M., Pall, G., et al. (2022). High-resolution single-cell atlas reveals diversity and plasticity of tissue-resident neutrophils in non-small cell lung cancer. Cancer Cell 40, 1503–1520 e1508. 10.1016/j.ccell.2022.10.008.

24. Zilionis, R., Engblom, C., Pfirschke, C., Savova, V., Zemmour, D., Saatcioglu, H.D., Krishnan, I., Maroni, G., Meyerovitz, C.V., Kerwin, C.M., et al. (2019). Single-Cell Transcriptomics of Human and Mouse Lung Cancers Reveals Conserved Myeloid Populations across Individuals and Species. Immunity 50, 1317–1334 e1310. 10.1016/j.immuni.2019.03.009.

25. Gentles, A.J., Newman, A.M., Liu, C.L., Bratman, S.V., Feng, W., Kim, D., Nair, V.S., Xu, Y., Khuong, A., Hoang, C.D., et al. (2015). The prognostic landscape of genes and infiltrating immune cells across human cancers. Nat Med 21, 938–945. 10.1038/nm.3909.

26. Faget, J., Peters, S., Quantin, X., Meylan, E., and Bonnefoy, N. (2021). Neutrophils in the era of immune checkpoint blockade. J Immunother Cancer 9. 10.1136/jitc-2020-002242.

27. Wei, X., Qian, W., Narasimhan, H., Chan, T., Liu, X., Arish, M., Young, S., Li, C., Cheon, I.S., Yu, Q., et al. (2025). Macrophage peroxisomes guide alveolar regeneration and limit SARS-CoV-2 tissue sequelae. Science 387, eadq2509. 10.1126/science.adq2509.

28. Wong, L.R., Zheng, J., Wilhelmsen, K., Li, K., Ortiz, M.E., Schnicker, N.J., Thurman, A., Pezzulo, A.A., Szachowicz, P.J., Li, P., et al. (2022). Eicosanoid signalling blockade protects middle-aged mice from severe COVID-19. Nature 605, 146–151. 10.1038/s41586-022-04630-3.

29. Li, C., Qian, W., Wei, X., Narasimhan, H., Wu, Y., Arish, M., Cheon, I.S., Tang, J., de Almeida Santos, G., Li, Y., et al. (2024). Comparative single-cell analysis reveals IFN-gamma as a driver of respiratory sequelae after acute COVID-19. Sci Transl Med 16, eadn0136. 10.1126/scitranslmed.adn0136.

30. Huang, S., Zhu, B., Cheon, I.S., Goplen, N.P., Jiang, L., Zhang, R., Peebles, R.S., Mack, M., Kaplan, M.H., Limper, A.H., and Sun, J. (2019). PPAR-gamma in Macrophages Limits Pulmonary Inflammation and Promotes Host Recovery following Respiratory Viral Infection. J Virol 93. 10.1128/JVI.00030-19.

31. Narasimhan, H., Cheon, I.S., Qian, W., Hu, S.S., Parimon, T., Li, C., Goplen, N., Wu, Y., Wei, X., Son, Y.M., et al. (2024). An aberrant immune-epithelial progenitor niche drives viral lung sequelae. Nature 634, 961–969. 10.1038/s41586-024-07926-8.

32. DuPage, M., Dooley, A.L., and Jacks, T. (2009). Conditional mouse lung cancer models using adenoviral or lentiviral delivery of Cre recombinase. Nat Protoc 4, 1064–1072. 10.1038/nprot.2009.95.

33. Miller, Y.E., Dwyer-Nield, L.D., Keith, R.L., Le, M., Franklin, W.A., and Malkinson, A.M. (2003). Induction of a high incidence of lung tumors in C57BL/6 mice with multiple ethyl carbamate injections. Cancer Lett 198, 139–144. 10.1016/s0304-3835(03)00309-4.

34. Pollard, A.J., and Bijker, E.M. (2021). A guide to vaccinology: from basic principles to new developments. Nat Rev Immunol 21, 83–100. 10.1038/s41577-020-00479-7.

35. Han, G., Sinjab, A., Rahal, Z., Lynch, A.M., Treekitkarnmongkol, W., Liu, Y., Serrano, A.G., Feng, J., Liang, K., Khan, K., et al. (2024). An atlas of epithelial cell states and plasticity in lung adenocarcinoma. Nature 627, 656–663. 10.1038/s41586-024-07113-9.

36. Marjanovic, N.D., Hofree, M., Chan, J.E., Canner, D., Wu, K., Trakala, M., Hartmann, G.G., Smith, O.C., Kim, J.Y., Evans, K.V., et al. (2020). Emergence of a High-Plasticity Cell State during Lung Cancer Evolution. Cancer Cell 38, 229–246 e213. 10.1016/j.ccell.2020.06.012.

37. de Visser, K.E., and Joyce, J.A. (2023). The evolving tumor microenvironment: From cancer initiation to metastatic outgrowth. Cancer Cell 41, 374–403. 10.1016/j.ccell.2023.02.016.

38. Cui, C., Wang, J., Fagerberg, E., Chen, P.M., Connolly, K.A., Damo, M., Cheung, J.F., Mao, T., Askari, A.S., Chen, S., et al. (2021). Neoantigen-driven B cell and CD4 T follicular helper cell collaboration promotes anti-tumor CD8 T cell responses. Cell 184, 6101–6118 e6113. 10.1016/j.cell.2021.11.007.

39. Engblom, C., Pfirschke, C., Zilionis, R., Da Silva Martins, J., Bos, S.A., Courties, G., Rickelt, S., Severe, N., Baryawno, N., Faget, J., et al. (2017). Osteoblasts remotely supply lung tumors with cancer-promoting SiglecF(high) neutrophils. Science 358. 10.1126/science.aal5081.

40. Pfirschke, C., Engblom, C., Gungabeesoon, J., Lin, Y., Rickelt, S., Zilionis, R., Messemaker, M., Siwicki, M., Gerhard, G.M., Kohl, A., et al. (2020). Tumor-Promoting Ly-6G(+) SiglecF(high) Cells Are Mature and Long-Lived Neutrophils. Cell Rep 32, 108164. 10.1016/j.celrep.2020.108164.

41. Gungabeesoon, J., Gort-Freitas, N.A., Kiss, M., Bolli, E., Messemaker, M., Siwicki, M., Hicham, M., Bill, R., Koch, P., Cianciaruso, C., et al. (2023). A neutrophil response linked to tumor control in immunotherapy. Cell 186, 1448–1464 e1420. 10.1016/j.cell.2023.02.032.

42. Ballesteros, I., Rubio-Ponce, A., Genua, M., Lusito, E., Kwok, I., Fernandez-Calvo, G., Khoyratty, T.E., van Grinsven, E., Gonzalez-Hernandez, S., Nicolas-Avila, J.A., et al. (2020). Co-option of Neutrophil Fates by Tissue Environments. Cell 183, 1282–1297 e1218. 10.1016/j.cell.2020.10.003.

43. Hajishengallis, G., Netea, M.G., and Chavakis, T. (2025). Trained immunity in chronic inflammatory diseases and cancer. Nat Rev Immunol. 10.1038/s41577-025-01132-x.

44. Aegerter, H., Kulikauskaite, J., Crotta, S., Patel, H., Kelly, G., Hessel, E.M., Mack, M., Beinke, S., and Wack, A. (2020). Influenza-induced monocyte-derived alveolar macrophages confer prolonged antibacterial protection. Nat Immunol 21, 145–157. 10.1038/s41590-019-0568-x.

45. Wang, T., Zhang, J., Wang, Y., Li, Y., Wang, L., Yu, Y., and Yao, Y. (2023). Influenza-trained mucosal-resident alveolar macrophages confer long-term antitumor immunity in the lungs. Nat Immunol 24, 423–438. 10.1038/s41590-023-01428-x.

46. Ryu, S., Shin, J.W., Kwon, S., Lee, J., Kim, Y.C., Bae, Y.S., Bae, Y.S., Kim, D.K., Kim, Y.S., Yang, S.H., and Kim, H.Y. (2022). Siglec-F-expressing neutrophils are essential for creating a profibrotic microenvironment in renal fibrosis. J Clin Invest 132. 10.1172/JCI156876.

47. Teo, J.M.N., Chen, Z., Chen, W., Tan, R.J.Y., Cao, Q., Chu, Y., Ma, D., Chen, L., Yu, H., Lam, K.H., et al. (2025). Tumor-associated neutrophils attenuate the immunosensitivity of hepatocellular carcinoma. J Exp Med 222. 10.1084/jem.20241442.

48. Kloosterman, D.J., and Akkari, L. (2023). Macrophages at the interface of the co-evolving cancer ecosystem. Cell 186, 1627–1651. 10.1016/j.cell.2023.02.020.

49. Arish, M., Qian, W., Narasimhan, H., and Sun, J. (2023). COVID-19 immunopathology: From acute diseases to chronic sequelae. J Med Virol 95, e28122. 10.1002/jmv.28122.

50. Han, X., Chen, L., Fan, Y., Alwalid, O., Jia, X., Zheng, Y., Liu, J., Li, Y., Cao, Y., Gu, J., et al. (2023). Longitudinal Assessment of Chest CT Findings and Pulmonary Function after COVID-19 Infection. Radiology 307, e222888. 10.1148/radiol.222888.

51. Cheon, I.S., Li, C., Son, Y.M., Goplen, N.P., Wu, Y., Cassmann, T., Wang, Z., Wei, X., Tang, J., Li, Y., et al. (2021). Immune signatures underlying post-acute COVID-19 lung sequelae. Sci Immunol 6, eabk1741. 10.1126/sciimmunol.abk1741.

52. Lopez-Otin, C., Pietrocola, F., Roiz-Valle, D., Galluzzi, L., and Kroemer, G. (2023). Meta-hallmarks of aging and cancer. Cell Metab 35, 12–35. 10.1016/j.cmet.2022.11.001.

53. Chia, S.B., Johnson, B.J., Hu, J., Vermeulen, R., Chadeau-Hyam, M., Guntoro, F., Montgomery, H., Boorgula, M.P., Sreekanth, V., Goodspeed, A., et al. (2024). Respiratory viral infection promotes the awakening and outgrowth of dormant metastatic breast cancer cells in lungs. Res Sq. 10.21203/rs.3.rs-4210090/v1.

54. Hill, W., Lim, E.L., Weeden, C.E., Lee, C., Augustine, M., Chen, K., Kuan, F.C., Marongiu, F., Evans, E.J., Jr., Moore, D.A., et al. (2023). Lung adenocarcinoma promotion by air pollutants. Nature 616, 159–167. 10.1038/s41586-023-05874-3.

55. England, F.J., Bordeu, I., Ng, M.E., Bang, J., Kim, B., Choi, J., Cardoso, E.C., Koo, B.K., Simons, B.D., and Lee, J.H. (2025). Sustained NF-kappaB activation allows mutant alveolar stem cells to co-opt a regeneration program for tumor initiation. Cell Stem Cell 32, 375–390 e379. 10.1016/j.stem.2025.01.011.

56. Maas, R.R., Soukup, K., Fournier, N., Massara, M., Galland, S., Kornete, M., Wischnewski, V., Lourenco, J., Croci, D., Alvarez-Prado, A.F., et al. (2023). The local microenvironment drives activation of neutrophils in human brain tumors. Cell 186, 4546–4566 e4527. 10.1016/j.cell.2023.08.043.

57. Barry, S.T., Gabrilovich, D.I., Sansom, O.J., Campbell, A.D., and Morton, J.P. (2023). Therapeutic targeting of tumour myeloid cells. Nat Rev Cancer 23, 216–237. 10.1038/s41568-022-00546-2.

58. Casanova-Acebes, M., Dalla, E., Leader, A.M., LeBerichel, J., Nikolic, J., Morales, B.M., Brown, M., Chang, C., Troncoso, L., Chen, S.T., et al. (2021). Tissue-resident macrophages provide a pro-tumorigenic niche to early NSCLC cells. Nature 595, 578–584. 10.1038/s41586-021-03651-8.

59. Martinez-Usatorre, A., Kadioglu, E., Boivin, G., Cianciaruso, C., Guichard, A., Torchia, B., Zangger, N., Nassiri, S., Keklikoglou, I., Schmittnaegel, M., et al. (2021). Overcoming microenvironmental resistance to PD-1 blockade in genetically engineered lung cancer models. Sci Transl Med 13. 10.1126/scitranslmed.abd1616.

60. Mantovani, A., Allavena, P., Marchesi, F., and Garlanda, C. (2022). Macrophages as tools and targets in cancer therapy. Nat Rev Drug Discov 21, 799–820. 10.1038/s41573-022-00520-5.

61. Jin, C., Lagoudas, G.K., Zhao, C., Bullman, S., Bhutkar, A., Hu, B., Ameh, S., Sandel, D., Liang, X.S., Mazzilli, S., et al. (2019). Commensal Microbiota Promote Lung Cancer Development via gammadelta T Cells. Cell 176, 998–1013 e1016. 10.1016/j.cell.2018.12.040.

62. Zhu, B., Wu, Y., Huang, S., Zhang, R., Son, Y.M., Li, C., Cheon, I.S., Gao, X., Wang, M., Chen, Y., et al. (2021). Uncoupling of macrophage inflammation from self-renewal modulates host recovery from respiratory viral infection. Immunity 54, 1200–1218 e1209. 10.1016/j.immuni.2021.04.001.

